# Integrative analysis of rare variants and pathway information shows convergent results between immune pathways, drug targets and epilepsy genes

**DOI:** 10.1101/410100

**Authors:** Hoang T. Nguyen, Amanda Dobbyn, Alexander W. Charney, Julien Bryois, April Kim, Whitney Mcfadden, Nathan G. Skene, Laura M. Huckins, Weiqing Wang, Douglas M Ruderfer, Xinyi Xu, Menachem Fromer, Shaun M Purcell, Kasper Lage, Matthijs Verhage, August B. Smit, Jens Hjerling-Leffler, Joseph D. Buxbaum, Dalila Pinto, Xin He, Patrick F Sullivan, Eli A. Stahl

## Abstract

Trio family and case-control studies of next-generation sequencing data have proven integral to understanding the contribution of rare inherited and *de novo* single-nucleotide variants to the genetic architecture of complex disease. Ideally, such studies should identify individual risk genes of moderate to large effect size to generate novel treatment hypotheses for further follow-up. However, due to insufficient power, gene set enrichment analyses have come to be relied upon for detecting differences between cases and controls, implicating sets of hundreds of genes rather than specific targets for further investigation. Here, we present a Bayesian statistical framework, termed gTADA, that integrates gene-set membership information with gene-level *de novo* and rare inherited case-control counts, to prioritize risk genes with excess rare variant burden within enriched gene sets. Applying gTADA to available whole-exome sequencing datasets for several neuropsychiatric conditions, we replicated previously reported gene set enrichments and identified novel risk genes. For epilepsy, gTADA prioritized 40 risk genes (posterior probabilities > 0.95), 6 of which replicate in an independent whole-genome sequencing study. In addition, 30/40 genes are novel genes. We found that epilepsy genes had high protein-protein interaction (PPI) network connectivity, and show specific expression during human brain development. Some of the top prioritized EPI genes were connected to a PPI subnetwork of immune genes and show specific expression in prenatal microglia. We also identified multiple enriched drug-target gene sets for EPI which included immunostimulants as well as known antiepileptics. Immune biology was supported specifically by case-control variants from familial epilepsies rather than do novo mutations in generalized encephalitic epilepsy.

## Introduction

*De novo* mutations (DNMs) have been successfully used to identify genes associated with neurodevelopmental disorders (NDDs) ^1-8^. Recently, additional risk genes have been reported by meta-analyzing DNMs and rare case-control (CC) variants, an approach that has been particularly successful for autism spectrum disorders (ASD) ^9,10^. For epilepsy (EPI), multiple associated genes have been identified through DN based studies ^4,5,11^, and in recent years, a number of EPI significant genes have also been identified through CC studies ^12,13^. We hypothesized that, as for ASD, additional significant EPI genes could be discovered through the integration of DN and CC data. EPI is a serious brain disorder which includes multiple subtypes. Studies of cases/controls and twins have shown that genetic components have played important roles in EPI ^14-16^. Some of EPI’s subtypes can be explained by single genes, but multiple subtypes might be caused by multiple genes ^15^. It is still challenging to develop specific drugs for this disorder. There have been multiple antiepileptic drugs used for EPI treatments; however, 20-30% of EPI patients have not been successful in controlling their seizures by using current medications ^17^. Identifying additional genes or gene sets might help better understand its etiology as well as better design drug targets for the disorder.

Due to the high polygenicity of NDDs, gene set (GS) tests have also been used to identify specific pathways relevant to disease etiology ^18-23^. A typical approach is that top significant genes are tested for enrichment in established sets and pathways. We here propose an alternative method that circumvents this issue by jointly modeling CC/DN variants and gene set information.

In this work, we introduce a method that tests gene-set enrichment directly from DN and rare CC data, and leverages enriched gene sets (eGSs) to prioritize risk genes. This approach allows genes to be prioritized if they are in enriched GSs/tissues, for a given strength of genetic evidence. The method can be used for discrete or continuous gene-set data and therefore can incorporate gene expression data to obtain additional significant genes based on tissue or cell-type expression information. It is a **g**eneralized framework of our extended **T**ransmission **A**nd **D**e novo **A**ssociation, gTADA. We apply gTADA to large DN and rare CC variant data, incorporating candidate and canonical gene sets, drug-target gene sets, and GTEx expression data in order to prioritize NDD and congenital heart disease (CHD) genes. With recent large rare CC data sets for EPI, we further analyze results for this disorder. We identify multiple significant EPI genes, and validate top genes in an independent data set. We provide further support for our significant genes through the analysis of expression data and protein-protein interaction networks.

## Results

We have developed the gTADA pipeline to prioritize risk genes for complex genetic disorders through integration of DN mutations, rare CC variants, and gene-set (GS) membership (Figure 1). The pipeline uses the **T**ransmission **A**nd **D**enovo **A**ssociation (TADA^9^ and extTADA^18^) framework to model and integrate DN and CC data, and combines gene-set information using a logistic regression model (Figure 1) in a **g**eneralized **TADA** ^9,18^ framework.

**Figure 1:**
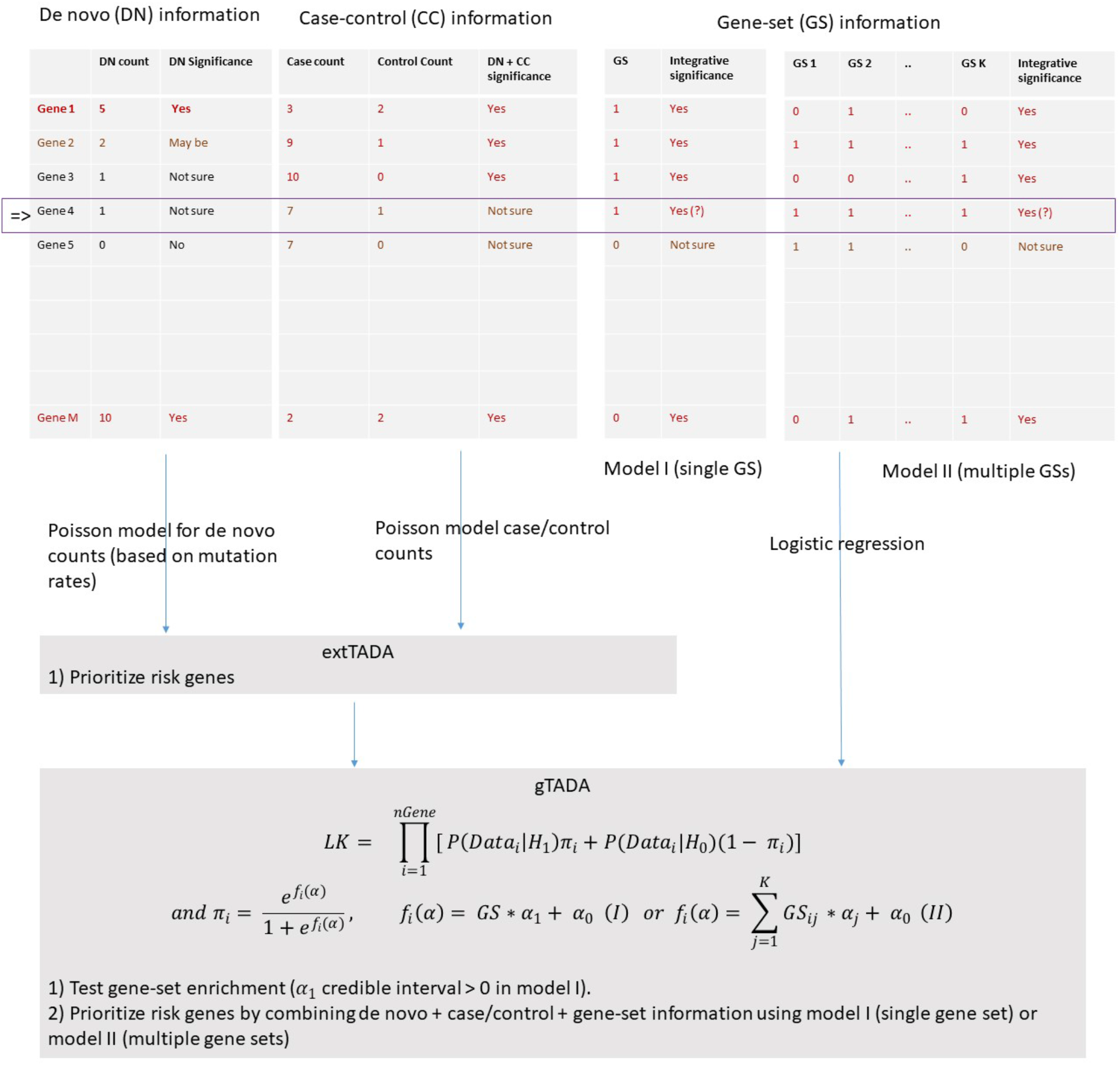
The framework of gTADA. The pipeline combines de novo (DN), case/control (CC) data (via variant counts of genes) and gene set (GS) information. It can test the enrichment of GS directly from the data (use α_1_ information from single-GS model), and prioritize risk genes using model I (single GS) or model II (multiple GSs). For example, Gene 4 might have a small posterior probability (PP) to be a risk gene because it does not have strong genetic information; however, the gene’s PP would be high when it is supported by GS information from eGSs.

To summarize, for each gene, all variants in a variant category are collapsed and considered as a single count (*x*). Table S1 presents the details of statistical models of the counts, their parameters, and the hyper parameters of DN and CC data. For each gene, gTADA compares two hypotheses: it is a risk gene (H_1_) and it is not a risk gene (H_0_). Similar to TADA^9^, our model assumes that rare variant counts in a risk gene are elevated by *γ* fold compared to chance expectation, and *γ* follows a gamma distribution: *γ* ~ *gamma* (*γ̄* * *β,β*) in which *γ̄* is the mean relative risk and *β* is the dispersion parameter of *γ*. For non-risk genes, *γ* = 1. We assume that there is a probability *π_i_* for the *i^th^* gene to be a risk gene. This *π_i_* is connected to a GS by *π_i_* = *e*^*f*_*i*_(*α*)^/(1 + *e*^*f*_*i*_(*α*)^), and; *f*_*i*_(*α*) = *α*_0_ + *GS_i_* * *α*_1_ where *GS_i_* is the value of the GS at the *i^th^* gene which can be 0/1 or a continuous value. This is in contrast to TADA and extTADA, in which *π_i_* is assumed to be the same across all genes. gTADA’s approach is more reasonable than previous approaches because genes should have different probabilities of being risk genes. The likelihood for the data at the i^th^ gene is *P*(*x*|*parameters*) = *P*(*x*|*H*_1_)*π_i_* + *P*(*x*|*H*_0_)(1 – *π_i_*). All parameters and hyper parameters of gene data were jointly estimated from the likelihood function across the all genes. As described in extTADA^24^, if variants are classified into different categories then similar statistical models are built separately for categories and their parameters are jointly estimated. The main model for testing GS enrichment and prioritizing significant genes was the single-GS model. We used a Markov Chain Monte Carlo (MCMC) method to sample parameters. Modes which were considered as the estimated values, and Bayesian credible intervals (CIs) of MCMC results were used in all the inferences. A GS was considered enriched if the lower boundary of its *α* CI was positive. One of the advantages of gTADA is that after learning gene sets, it can use that knowledge to increase the power of finding risk genes, because genes in the enriched GS will have higher prior probabilities ^24^. We used posterior probabilities (PPs) to prioritize risk genes with 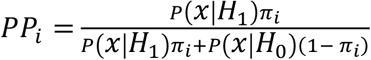 for the *i^th^* gene. After testing multiple gene sets separately, we reported genes with PP>0.95 with any gene set as significant gTADA genes, and also conducted follow-up analyses on significant and suggestive genes with PP>0.8 with any gene set. We conducted simulation analyses based on real data, to show the effect of taking the union of results across gene sets.

We simulated different DN/CC data using genetic parameters from previous ASD studies ^9,18^. Enriched gene sets (eGSs) were simulated using the results of known enriched gene sets for ASD ^18^ (see Methods), and non-enriched gene sets were simulated by randomly choosing genes. Different trio numbers were used in the simulation process ranging from 1,000 to 50,000. Genetic parameters of simulated data are presented in Table S2 (See Methods). We tested simulated data for single gene set and multiple gene set analyses (Supplementary Information). For single-GS models, the number of risk genes identified increased when eGSs were used (Figure S1). In addition, the Type I error of calling a GS enriched was well calibrated (Table S3). For multiple-GS models, the number of risk genes increased when eGSs numbers increased; however, we observed higher rates of false positive risk genes particularly with small sample sizes (Figure S2). For this reason, we focused our analyses on single gene set models, and combined results across single gene set models.

We applied gTADA to available rare variant data of four NDDs and CHD to prioritize genes for these disorders (Figure 2). In summary, this data included 4293, 1012, 1213, 5122 and 356 trios of DD, ID, CHD, ASD and EPI respectively; plus 4058 ASD and 5704 EPI case/control data (see Methods Data). These data were annotated and divided into different categories by using the approach of Nguyen, et al. ^18^. We used loss-of-function (LoF) and missense damaging (MiD) categories of these annotations. For EPI case/control, we only used count data from Epi K. consortium and Epilepsy Phenome/Genome Project ^13^ which were annotated by the authors (details in the Method). GSs were called enriched if their 95% credible intervals were larger than zero. GSs were further called significantly enriched (seGS) if their Benjamini and Hochberg ^25^ adjusted p value (pBH) was < 0.05. To identify significant genes for each eGS, we set a stringent maximum PP (PP_max_) threshold of 0.95. We also examined the properties of prioritized genes having PP_max_ > 0.8.

**Figure 2:**
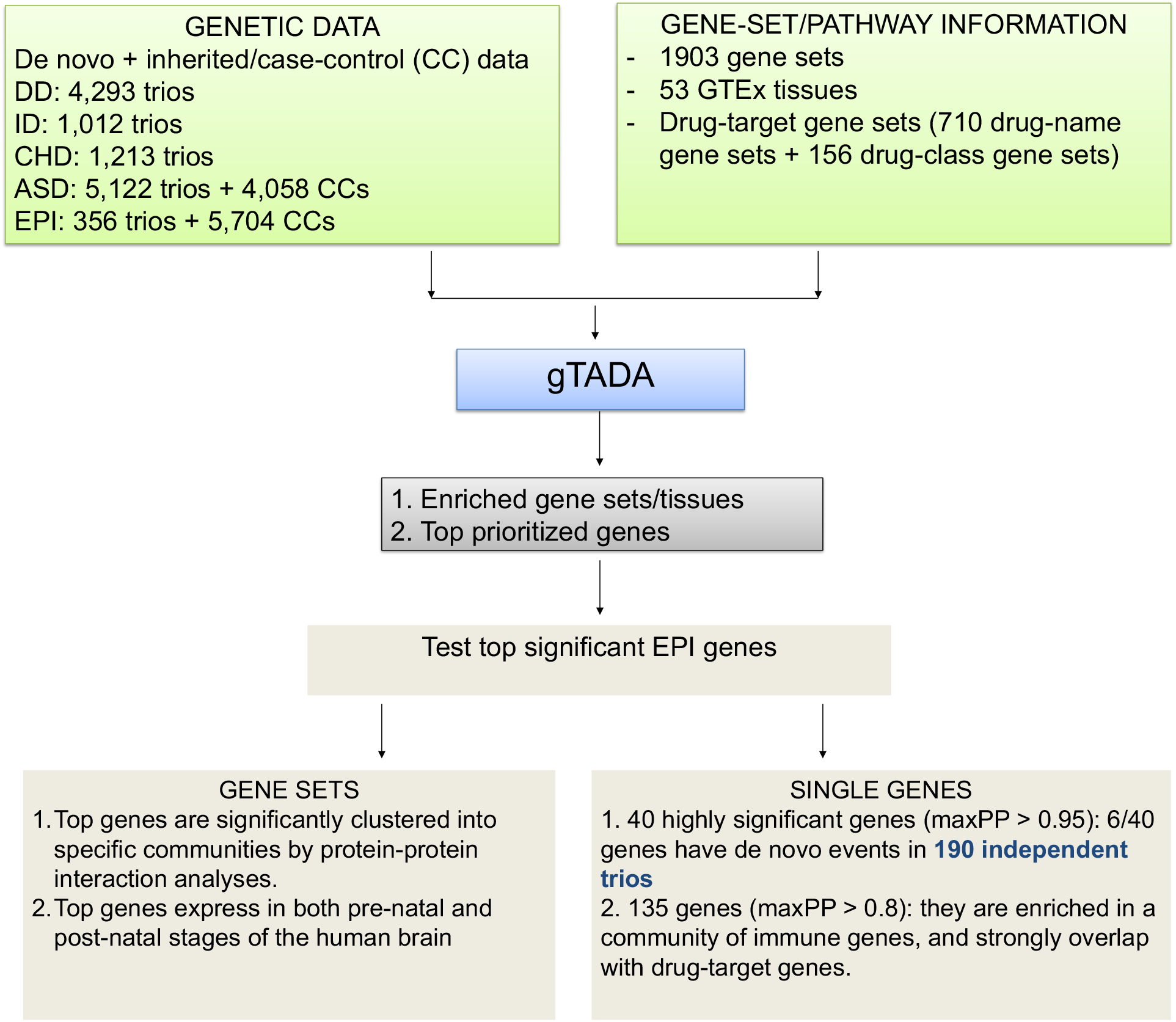
Data analyses in the study. Four neurodevelopmental disorders (NDDs) and congenital heart disease (CHD) are analyzed. Results of epilepsy (EPI) are validated by using different methods and an independent data set.

We tested 1,903 GSs used in our previous study ^18^, including 186 candidate and 1,717 gene sets with 100 to 4,955 genes from MSigDB ^26^ and the Gene Ontology data base ^27^ (Table S4, Table 1). gTADA identified multiple eGSs for all disorders (Table 1). All gTADA GS enrichment results are presented in Table S5. Overall, CHD, ASD, ID and DD had the highest overlapping seGSs (132 GSs, Figure S3). The top seGS of each disorder replicated previous results ^18,28^. gTADA was able to re-call >89% enriched gene sets reported by our previous results (Supplementary Information, Figure S4). To better understand the performance of gTADA on each eGS individually, we chose top 20 eGSs from each disorder based on significance, and compared the significant gene-count results of gTADA and extTADA using a threshold PP>0.95. ASD, DD and ID gained more significant genes than CHD and EPI when GSs were used (Figure 3). In addition, EPI had more small eGSs than the four other disorders.

**Figure 3:**
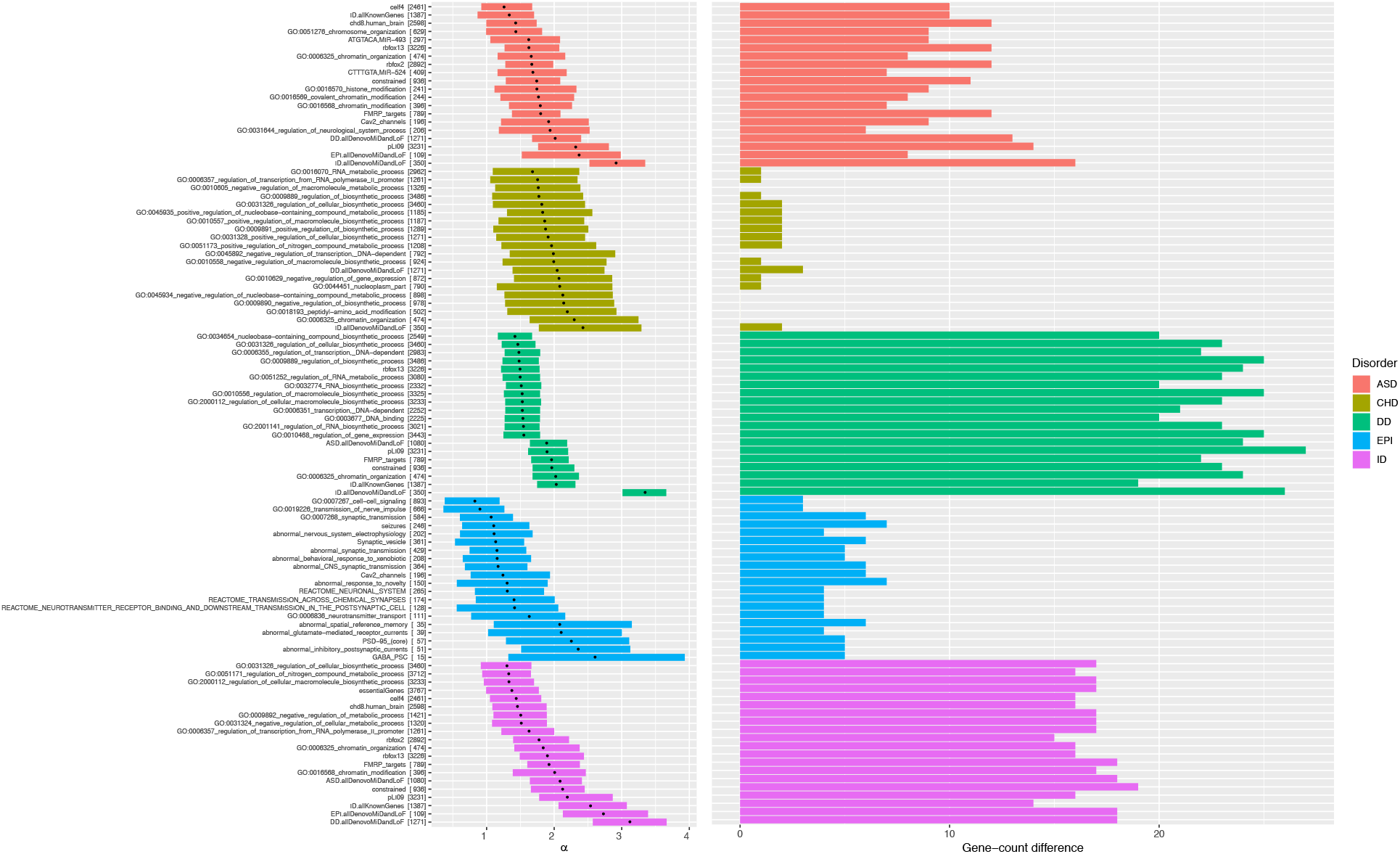
Top enriched gene sets (eGSs) of disorders. These are top eGSs of the analyzed disorders (based on α_1_’s significance). Y-axes are names of the eGSs and their sizes (e.g, GO:0016568 has 396 genes). The left picture shows α_1_’s credible intervals and modes of eGSs. The right picture describes the differences in gene counts (posterior probabilities > 0.95) between using GSs and not using GSs.

**Table 1:** The number of prioritized genes for all disorders. n GS and eGS are the number of tested GSs and the number of enriched/significant GSs (lower CIs > 0) respectively. For each disorder, we did not use its own DNM and known gene sets to avoid inflating results. seGS is the number of tested GSs whose lower CIs are > 0 and adjusted p values are < 0.05. For each column with maximum posterior probability (PP) > a threshold, the number in each cell is the number of prioritized genes.

| Disorder | Results: number of tested/significant gene sets/tissues |  |  |  |  |  |  |  |  |  |  |  | Results: number of prioritized genes |  |  |  |  |  |  |  |
| --- | --- | --- | --- | --- | --- | --- | --- | --- | --- | --- | --- | --- | --- | --- | --- | --- | --- | --- | --- | --- |
|  | Candidate GS |  |  | GTEx tissue |  |  | Drug-target GS |  |  | Drug-target (class) GS |  |  | Candidate GS |  | GTEx tissue |  | Drug-name GS |  | Drug-class GS |  |
|  | n GS | eGS | seGS | n GS | eGS | seGS | n GS | eGS | seGS | n GS | eGS | seGS | PP<sub>max</sub>>0.95 | PP<sub>max</sub>>0.8 | PP<sub>max</sub>>0.95 | PP<sub>max</sub>>0.8 | PP<sub>max</sub>>0.95 | PP<sub>max</sub>>0.8 | PP<sub>max</sub>>0.95 | PP<sub>max</sub>>0.8 |
| ASD | 1901 | 381 | 338 | 53 | 29 | 28 | 710 | 31 | 0 | 156 | 3 | 0 | 63 | 191 | 33 | 59 | 35 | 63 | 31 | 57 |
| ID | 1901 | 495 | 485 | 53 | 52 | 52 | 710 | 36 | 2 | 156 | 8 | 4 | 59 | 177 | 45 | 74 | 47 | 63 | 45 | 60 |
| DD | 1901 | 686 | 679 | 53 | 53 | 53 | 710 | 39 | 3 | 156 | 4 | 0 | 167 | 288 | 129 | 198 | 115 | 159 | 114 | 152 |
| EPI | **1901** | **108** | **50** | **53** | **9** | **6** | **710** | **88** | **67** | **156** | **18** | **13** | **40** | **135** | **26** | **82** | **35** | **100** | **33** | **95** |
| CHD | 1902 | 280 | 241 | 53 | 50 | 50 | 710 | 0 | 0 | 156 | 2 | 0 | 12 | 101 | 6 | 16 | 0 | 0 | 5 | 15 |

We combined results from all eGSs to prioritize risk genes for each disorder. Based on PP_max_>0.95, DD had the highest number of genes (167) followed by ASD, ID and EPI (63, 59 and 40 respectively) while CHD had only 12 prioritized genes (Table 1). All prioritized genes are presented in Table S6. One gene, STXBP1, was observed across all four NDDs with PP_max_ > 0.95.

In addition, 18 genes were identified (PP_max_ > 0.95) in at least three disorders (Table S6). The results of gTADA were from combining multiple single-GS analyses; therefore, we tested the observed false discovery rates and saw that PP_max_>0.95 was nearly equivalent to FDR<0.1 (Figure S5, Supplementary Information).

We also applied gTADA to expression data of 53 tissues from GTEx Consortium ^29^. Only 6 tissues were enriched for EPI while 28 issues were enriched for ASD (Table 1). Interestingly, >= 50 tissues showed enrichment for ID, DD and CHD. ID and DD data were very well powered, while CHD risk genes may be highly expressed across multiple tissues. All enrichment results are presented in Table S7 and in Figure S6. The risk-gene numbers from enriched tissues were not as high as results from candidate GSs. One possible reason was that the estimated *α*_1_ values of GTEx were not high (less than 1, Table S7) because we used continuous values for all genes. There were 11 genes with PP > 0.95 in at least three disorders. All these 11 were among the 15 genes identified with candidate and canonical GSs above (Table S8).

We also tested drug target GSs described in Ruderfer, et al. ^30^. Briefly, drug target genes were predicted by using the Similarity Ensemble Approach (SEA) ^31^ on data from DrugBank version 4.1 ^32^ and ChEMBL-14. We tested 710 and 156 drug and drug-class GSs based on the Anatomical Therapeutic Chemical (ATC) classification system Level 3 and Level 5 respectively. EPI had the highest number of significantly enriched drug GSs, followed by ID and DD (67, 3 and 2 seGSs respectively; Table 1 and S9). There were 13 and 4 significantly enriched drug-class GSs for EPI and ID respectively (Table 1 and S10). There were some eGSs for other disorders, but none were significant after adjusting for multiple tests. Two drug classes were observed in the enrichment results of both EPI and ID: ANTIEPILEPTICS (pBH = 2.1×10e−3 and 0.015 respectively) and ANTIPROPULSIVES (pBH = 0.01 for both disorders) (Table S10). Interestingly, some immune drug target GSs were significant or nominally significant for EPI: the drug nabumetone (pBH = 0.02; a member of drug-class ANTIINFLAMMATORY AND ANTIRHEUMATIC PRODUCTS, NON-STEROIDS), and the drug-class GSs OTHER DERMATOLOGICAL PREPARATIONS (pBH = 1.1e-3), IMMUNOSTIMULANTS (pBH = 0.003), ANTIINFLAMMATORY AGENTS (pBH = 0.06). To test whether the enrichment of immune drug target GSs was driven by their overlap with ANTIEPILEPTICS, we re-ran gTADA on the drug-class GSs after removing overlapping genes with the ANTIEPILEPTICS GS. The drug-class IMMUNOSTIMULANTS remained significant (pBH = 0.013). Prioritized risk genes using enriched drug target GSs are shown in Tables 1, S11, S12.

We focus on EPI because this disorder had DN and CC data, including recent rare CC variant studies ^12,13^. In addition, multiple EPI genes were prioritized by gTADA. gTADA results without GSs provide an estimated proportion of risk genes of 4.9% (Table S13), higher than the proportion estimated in Nguyen, et al. ^18^, however Nguyen et al., 2017 only used DN data. Based on this proportion, the mean DN RRs (estimated *γ̄*) were >15 (16 and 18 for MiD and LoF mutations respectively). The mean RRs of the three CC samples were > 4 (Supplementary Information). Details of the analyses are in Section 1.2 of Supplementary Information, Figure S7 and Table S14.

We sought to validate gTADA identified EPI risk genes, focusing on the results from 1,903 candidate/canonical GSs, from which higher numbers of significant genes were obtained, and which included all genes prioritized using GTEx and most genes prioritized using the drug-target GSs (Figure S8). gTADA prioritized 40 genes with PPmax>0.95 from 108 eGSs (Table S6), 30 of which (*ATP8B1, C5orf42, CACNA1B, CEP89, COPB1, CSNK1E, DRC1, EHD4, FGFR1OP, FURIN, GABBR2, GIGYF1, GPAM, GPR87, GRIA4, HSD17B4, KDM6B, KEAP1, NFATC3, NRXN2, PHTF1, PMPCA, SAMD9L, SCYL1, SLC10A1, SLC8A2, SLC9A2, TBCK, TRMT1L, TYRO3*) were not in the list of known EPI genes. One gene (*GABBR2*) was identified in our recent analysis of only DN data ^18^. ExAC LoF-intolerance pLI information ^33^ was available for 29 of the 30 novel genes. 12/29 genes (*CACNA1B, COPB1, CSNK1E, FURIN, GABBR2, GIGYF1, GRIA4, KDM6B, NFATC3, NRXN2, TRMT1L, TYRO3*) were highly intolerant genes (pLI > 0.9). Interestingly, 13/30 genes (*ATP8B1, C5orf42, CEP89, DRC1, EHD4, GPAM, HSD17B4, PHTF1, PMPCA, SAMD9L, SCYL1, SLC10A1, TBCK*) had pLI < 0.1 and were not known missense constrained genes ^34^. We investigated these genes and saw that the significant signal of these 11 genes was from CC data. *C5orf42, HSD17B4, PMPCA, SAMD9L, SCYL1* and *TBCK* were also reported in NDD studies ^35-40^. The Epi K. consortium and Epilepsy Phenome/Genome Project ^13^ used the same CC data set as our current study and reported 7 significant autosomal genes (*DEPDC5, GABRG2, GRIN2A, KCNQ2, LGI1, PCDH19, SCN1A*); all seven had gTADA PP_max_ > 0.9, and 6/7 (except *GRIN2A)* had PP_max_ > 0.95. We also saw that the majority of the 40 genes which had PP_max_>0.95 were inside eGSs (Figure 4).

**Figure 4:**
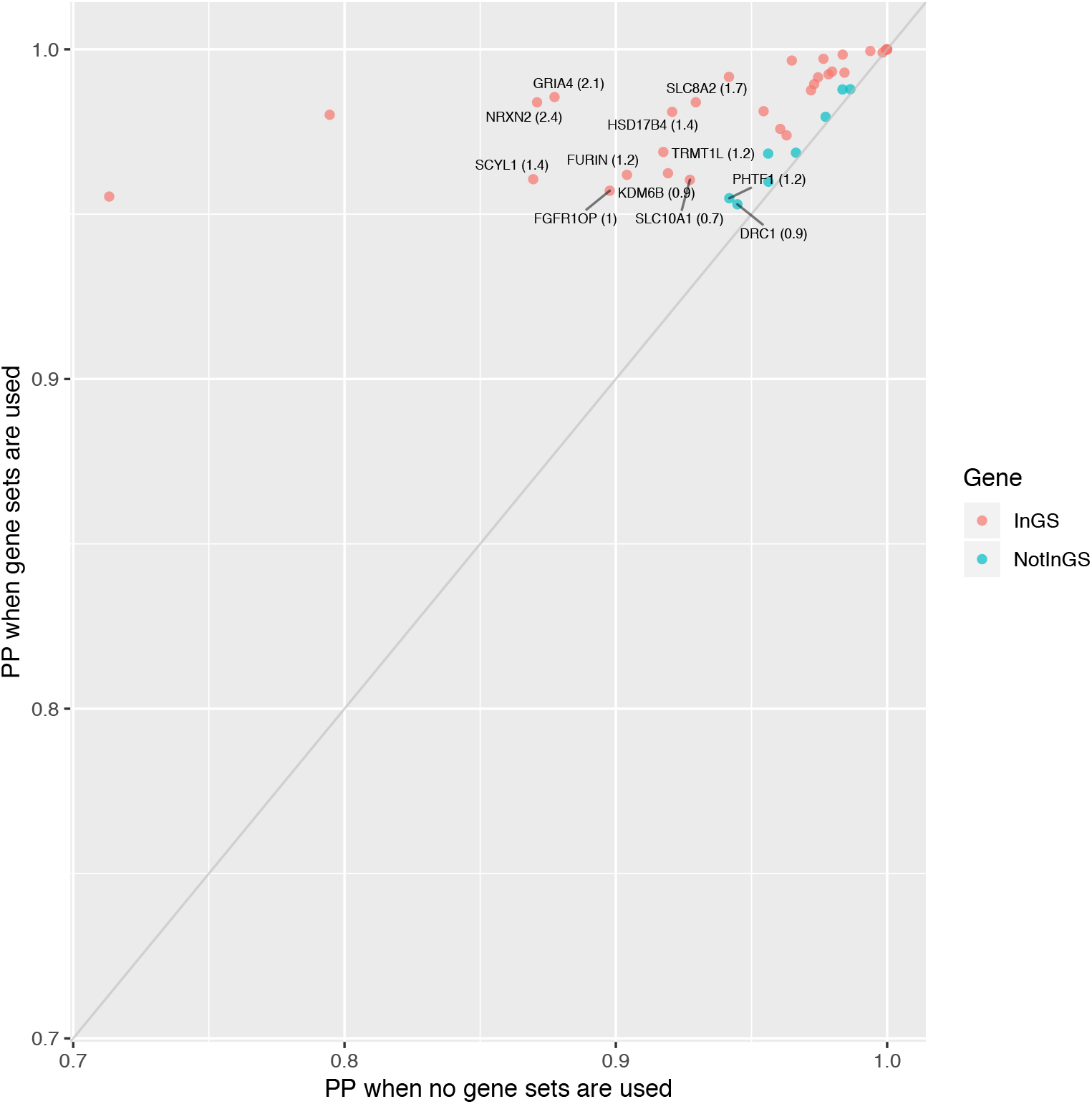
Comparing posterior probabilities (PPs) for top epilepsy (EPI) genes (PP > 0.95). The x-axis shows the PPs when no gene sets are used while the y-axis shows the PPs when enriched GSs are used. Points with gene names describe novel risk genes whose PPs are less than 0.95 if no GSs are used. Genes in the red color are inside enriched GSs while genes in the blue color are not inside enriched GSs.

Recently, Hamdan, et al. ^41^ sequenced the whole genomes of 197 trios with developmental and epileptic encephalopathy (DEE). From the 40 genes identified by gTADA using candidate/canonical GSs, 6 genes (*CSNK1E, GABBR2, GABRG2, GNAO1, KCNQ2, SCN1A*) had DNMs in the 197 trios (p value for this overlap < 5.9e-5). Interestingly, *SCNA1* had 6 DNMs and *GNAO1* had 2. Two of the 30 novel gTADA genes, *GABBR2* and *CSNK1E*, had one nonsynonymous DNM each. The gene *GABRB2* was reported as a significant risk gene for DEE by Hamdan, et al. ^41^ because it was in a *de novo* copy-number variant (CNV) duplication in 6 probands.

We analyzed the protein-protein interaction (PPI) network connectivity of 135 top gTADA EPI genes (PP_max_ > 0.8) using GeNets ^42^. We found that 100/135 genes and 57 direct connection candidate genes were well connected in five communities (overall and community connectivity p values < 2e-3, Figure 5A). The communities showed enrichment for multiple canonical pathways (Table S15): ion channel transport, neurotransmitter receptor binding, GABA receptor activation, ligand gated ion channel transport (Community 2); JAK-STAT signaling pathway, regulation of IFNA signaling, RIG-I-like receptor signaling pathway, interferon alpha/beta signaling, and autoimmune thyroid disease (Community 4).

**Figure 5:**
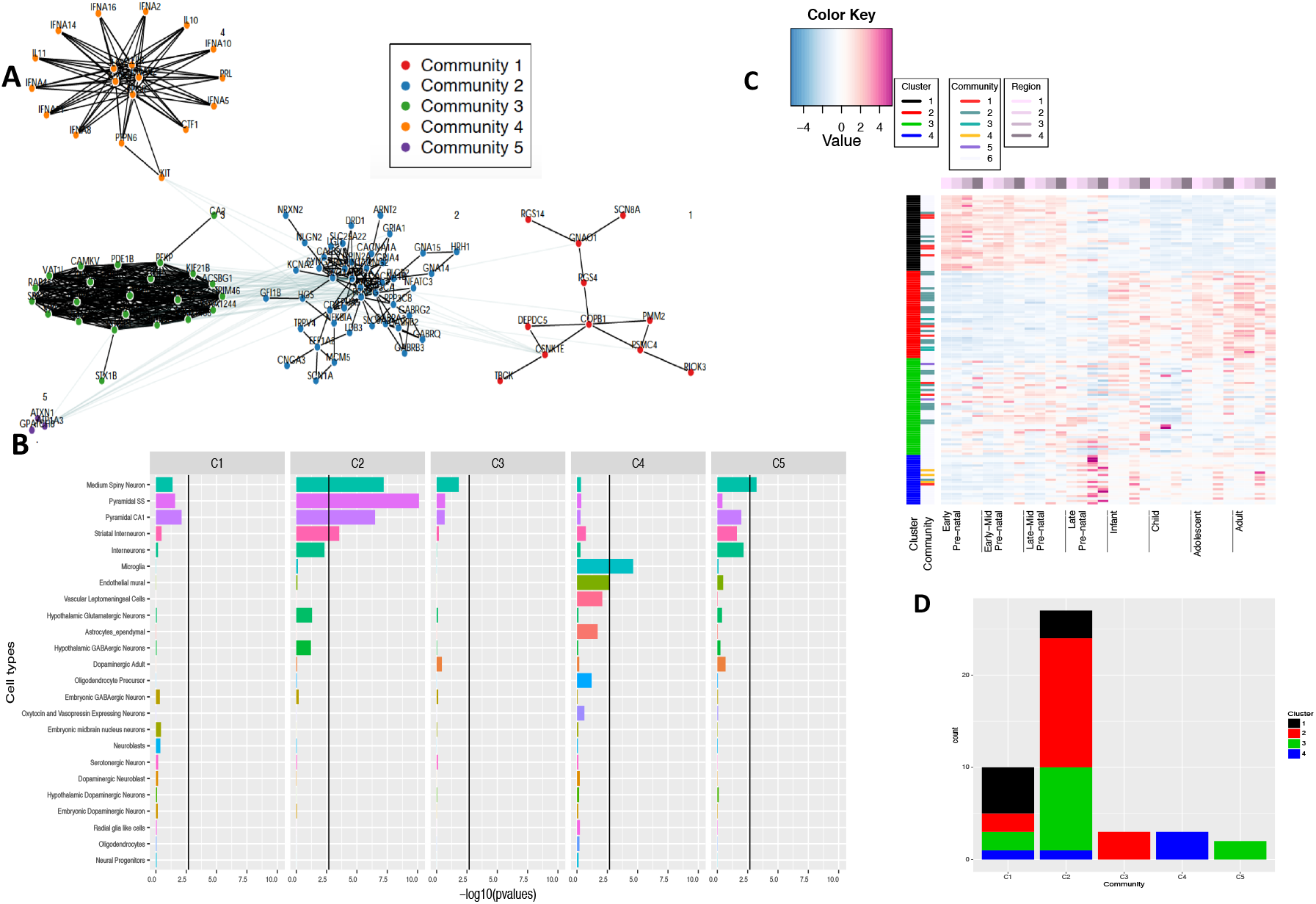
Results of the top prioritized EPI genes. These are genes whose maximum posterior probabilities are larger than 0.8. A: Protein-protein interaction (PPI) analysis for these genes; B: enrichment results of different mouse cell types using single-cell RNA data for Communities; C: spatiotemporal gene expressions across the genes in four regions (frontal cortex, temporal and parietal cortical regions, sensory-motor regions, and subcortical regions) of the human brain; D: gene counts in spatiotemporal brain expression clusters from C for PPI Communities from A.

The InWeb/GeNets^42,43^ PPI data are highly curated but nevertheless include many interactions from high-throughput in vitro assays that are false positive in the sense of in vivo biological function. To assess the influence of incorporating high throughput PPI data, we also used the STRING database ^44^ to test for physical interactions among EPI genes from gTADA and from our GeNets communities. Using only sources with experimental evidence, PPI connectivity was significant among the 135 gTADA EPI genes (13 observed edges versus 7 expected edges, p = 0.0248), and was strongly significant for Communities 2 and 4 (p < 4.33e-10). Interaction signals were weaker for other communities (p = 0.02, 0.07 and no interactions for Community 1, 3 and 5 respectively).

We tested EPI gene PPI communities for specific enrichment in our recent high-depth mouse brain single cell RNA sequencing (scRNAseq) data ^45^, which included fetal cell types (Figure 5B). We saw enrichment pyramidal CA1, SS neuronal expression in Communities 2 and 5, similar to results we recently reported for four NDDs ^18^. Interestingly, Community 4 gene expression was enriched in microglia cells (p = 2.34e-5). Endothelial mural, vascular leptomeningeal, astrocytes_ependymal cells were also enriched in Community 4 but not as strongly as microglia. Similarly, scRNAseq cell type enrichments were seen for these communities using only gTADA EPI genes and not GeNets candidate genes (Figure S9). Microglia were enriched for Community 4 (p = 1.5e-3), followed by endothelial mural cells (p = 0.02). Community 4 included three gTADA genes *IFNAR2, LEPR, PTPN6; PTPN6 and IFNAR2* were strongly specifically expressed in microglia (Figure S10). Observing genes with PP_max_>0.8 from drug-target based gTADA, *IFNAR2* was prioritized as a top gene in multiple drug-target GSs with the strongest PP from the drug class IMMUNOSTIMULANTS (PP = 0.99) while *PTPN6* was also prioritized by the drug-name GS mebeverine (PP = 0.85, in class of drugs for functional gastrointestinal disorders). In available human brain scRNAseq data sets ^46,47^ gTADA EPI genes were enriched in neuronal and GABAergic cell types (Figure S11), with the strongest enrichments observed for Community 2. These results were similar to the results of mouse cell types in Figure 5. Community 4 showed positive but non-significant microglial enrichment in human brain scRNAseq cell types (Figure S11). In human fetal brain scRNAseq data ^48^, PTPN6 was also present in clusters of immune and microglia cell types.

We next examined our prioritized EPI genes in the BrainSpan^49^ spatiotemporal brain gene expression data. The EPI prioritized genes showed expression during all developmental stages of the human brain (Figure 5C). Hierarchical clustering of EPI gene spatiotemporal brain expression identified expression patterns that were largely prenatal (black); postnatal (red); prenatal, infant, and postnatal in the cerebellum (green); and late prenatal, and postnatal in striatal regions (blue). In contrast, DD, ID, and CHD genes were more strongly expressed in prenatal stages (Figure S12). Spatiotemporal expression correlated with PPI communities (p = 0.0007153, Figure 5D); PPI communities 1 and 2 had higher proportions of genes with specific prenatal and postnatal expression, respectively, while Communities 3, 4 and 5 had only genes from single expression clusters (red, green and blue respectively). gTADA genes in the immune PPI community (*IFNAR2, LEPR, PTPN6*) were strongly expressed in late prenatal stages particularly in subcortical regions, and postnatally in the striatum (Figure 5C, S13).

A number of gene sets from the drug-target data were highly enriched in the EPI data, driven especially by GABA receptor genes. The genes *GABRG2, GABRA5, GABRA1* were all present in at least 45 of the 67, and the voltage-gated sodium channel genes *SCN1A, SCN8A* and *SCN2A* were present in at least 32, significantly enriched GSs (Figure 6).

**Figure 6:**
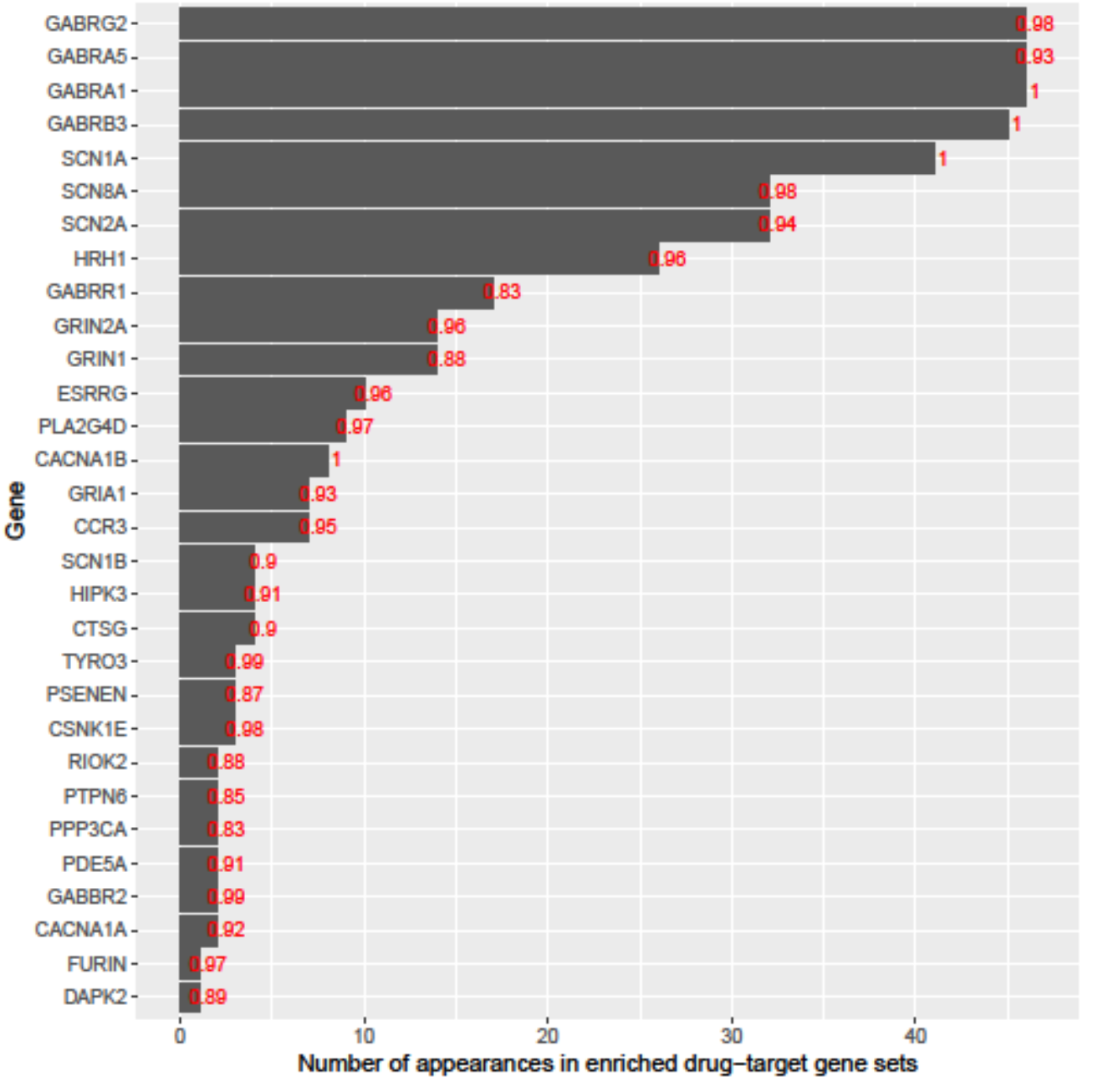
The number of appearances of genes in enriched drug-target gene sets. For each gene, the number in red is the highest gTADA posterior probability (PP_max_) of the gene.

To see whether the DN or CC signal was specific for these clusters, we counted all the DN mutations and calculated CC ratios for all PPI communities, spatiotemporal expression clusters and significantly enriched drug target gene sets. The DN signals of Communities 1, 2 and 3 were much stronger than those of other Communities (ratios of observed and expected DN counts > 80, which were highly larger than the meanRRs of DN signals). Regarding gene-expression clusters, surprisingly, multiple *de novo* mutations were observed in postnatal genes (Table S16).

For enriched drug-class GSs, both DN and CC data were enriched for these GSs, but DN signals were much higher than CC signals for the majority of GSs. In addition, the main signals of CC data for these enriched GSs were from familial non-acquired focal epilepsy (Table S16). Interestingly, very different from other GSs, IMMUNOSTIMULANTS was enriched because of only CC signals, and there were no GABA receptor genes in this GS; hence this GS remained significant for EPI after removing genes overlapping with the ANTIEPILEPTICS drug class targets.

## Discussion

We have presented a pipeline (gTADA) that incorporates *de novo* mutations (DNMs), rare inherited/case-control (CC) variants and pathway/gene-set/expression information to prioritize disease genes. This pipeline is based on our previous work, extTADA ^18^, but gTADA is a generalized framework of extTADA. gTADA can be extTADA if the gene-set information is not used. Recently, methods which use annotation/expression information to impute more risk genes have been actively developed for common variants ^50-52^. These methods have been successfully used to prioritize risk genes, and elucidate biological pathways for schizophrenia, bipolar ^18,53^ and breast cancer ^54^. gTADA might be the first tool using this approach for rare variants. There are many benefits of this approach ^50^. First, it can identify significant genes which might be missed by using typical genetic-data based methods. Second, significant genes can assist in understanding the structure of enriched gene sets. Another advantage of gTADA is that the package can test gene set enrichment directly from data. This enrichment test has been shown more powerful than other ways in the analyses of ChiP-Seq data sets ^55^. We hope that gTADA will be helpful in rare-variant based studies. The code is available online on https://github.com/hoangtn/gTADA.

We used gTADA to identify enriched tissue/gene sets (GSs) (from candidate GSs, drug-target GSs and GTEx tissues); and to prioritize genes for neurodevelopmental disorders (NDDs) and congenital heart disease (CHD). We saw that six human brain-region tissues and multiple candidate GS were enriched across NDDs and CHDs (Table 1). Interestingly, multiple drug target GSs were enriched in EPI, but just a few were enriched in DD and ID, and there were no enriched GS for ASD and CHD. In analyses leveraging the enriched GSs, multiple significant genes were identified for all these disorders (Table 1).

We analyzed EPI results in depth because new rare CC data sets were analyzed, and multiple novel EPI rick genes were identified. By combining the results from multiple gene sets, there were 40 genes with maximum posterior probabilities (PP_max_) > 0.95, corresponding to FDR<0.1. Thirty of our 40 EPI genes were not in the list of known EPI genes. Two of the 30 genes had *de novo* events in a new trio data set, and some of the 30 genes have been reported in other studies of neurodevelopmental disorders. This number of genes was much higher than analyses with only DN or only CC. The number of predicted risk genes of these EPI combined data sets (~ 950) was higher than that of the DN-only based genetic architecture ^18^. The EPI CC data represent three EPI types, familial non-acquired focal epilepsy (familial NAFE), familial genetic generalized epilepsy (familial GGE), and sporadic non-acquired focal epilepsy (NAFE), whereas the DN data are from epileptic encephalopathies (EE); combining heterogeneous DN and CC data could contribute to larger estimated proportion of risk genes. In a recent case/control study ^56^, risk genes were observed for familial non-acquired focal and familial genetic generalized epilepsies. However, in this study, gTADA estimated mean relative risks (RRs) were nearly equal for all three CC population samples: familial non-acquired focal epilepsy, familial genetic generalized epilepsy, and sporadic non-acquired focal epilepsy (Supplementary Information, Table S14). The top gTADA genes had higher differences in the variant counts between cases and controls than the other well-known gene sets, including known EPI genes and FMRP targets (Table S14). Therefore, this result suggests that sporadic non-acquired focal epilepsy top risk genes are the same as the two familial EPI types. The top drug target gene sets were supported by DN and CC data, with some genes occurring in many enriched GSs: *GABRG2, GABRA5, GABRA1, GABRB3, SCN1A* (Figure 6). These genes have been discussed as potential drug targets specific for EPI as well as other neurodevelopmental disorders ^57-59^. Further studies focusing on deeply understanding genetic variants in these genes could help better design drug targets for EPI.

In our current study, we observed that EPI genes with PP_max_ > 0.8 were well connected in five communities by a protein-protein interaction (PPI) network analysis. These genes showed expression in different developmental stages of the human brain (Figure 5). Interestingly, Community 4 from the PPI network analysis was enriched in immune pathways. Genes from this community were strongly expressed in the late prenatal stages of the human brain. In addition, based on scRNAseq, we saw that microglia cells were strongly enriched in this community but not in other communities. We note that the PPI results here rely on interaction data from high-throughput in vitro assays ^42,43^. The selection of members of Community 4 is based on these data. To verify these results, we used the STRING database with only edges with experimental evidence and still saw significant interactions between members inside Community 4. In addition, the results of Community 4 are supported by the enrichment of immune drug-target GSs. The relationship between epilepsy and the immune system as well as inflammatory pathways has been discussed elsewhere ^60-64^. Future studies exploring more on this connection could be beneficial in understanding the etiology of epilepsy.

While this study uses a novel approach to integrate different types of genomic data, it does have some limitations. First, gTADA partly relies on reference data sets (e.g., gene sets, tissues, Figure 3, Table 1). Many candidate gene sets may also be imperfect. For example, *STXBP1* is a well known presynaptic gene ^65,66^, but occurs in the significantly enriched PSD_Bayes2011 ^67^ gene set. However, gTADA is a model-based analysis of DN and CC variant data; therefore, the top prioritized genes are generally supported by the DN and CC data, not solely from reference data sets (Figure S8). In simulations when large GSs or multiple GSs are used, rates of false positive gene identification increase. One obvious reason is that if a GS size is larger than the number of risk genes, imputed genes outside the range of risk genes would be called false positive genes. However, for real data, large enriched gene sets might help in identifying more novel risk genes (Figure 3). For the current model with multiple GSs, we must calibrate an increased PP threshold for accurate control of FDR (Figure S1, S2). For example, we used a threshold PP_max_>0.95 to obtain top EPI genes. Based on simulation data from EPI genetic parameters, the prioritized genes should have FDRs < 0.1 (Figure S5). Using the same PP_max_ threshold, we also saw that FDRs increased quickly when few gene sets were added; however, FDRs slightly increased when more gene sets were added (Figure S5). This might be because the enriched GSs overlap with each other. As a result, more significant genes are not identified when additional GSs are added. Therefore, FDRs do not change much after adding a number of GSs. Simulations show (Figure S1) and as discussed by Nguyen, et al. ^18^, larger sample sizes and improved modeling approaches can help to address these weaknesses in future studies. gTADA as well as its previous pipelines ^9,18^ model variant-count data using statistical distributions (e.g, Poisson distribution for rare variants); therefore, count data should follow these distributions to obtain optimal results. Finally, the top prioritized EPI genes here are based on meta-analyzing multiple population samples and types of EPI; generally, as with any meta-analysis approach, heterogeneity should be assessed in the results. In this case, DN and multiple CC datasets support top EPI gene sets and risk genes, and many prioritized EPI genes were reported for other neurodevelopmental disorders, suggesting that the results point to convergent dysfunctions across EPI types and NDDs.

## Methods and data

### Data

#### Gene-set data

We used 1903 gene sets curated by Nguyen, et al. ^18^. These included 186 known gene sets with prior evidence of involvement in ASD and SCZ, and 1717 gene sets whose lengths were between 100 and 4995 genes from different databases: the Gene Ontology database ^68^, KEGG, and REACTOME, and the C3 motif gene sets from the Molecular Signatures Database (MSigDB) ^69^. The information of these gene sets was presented in Table S2 of Nguyen et al. 2017 and was also summarized in Table S4 in this study.

Drug-target gene sets were processed and classified as Ruderfer, et al. ^70^. Briefly, drugs were classified according to the level of the Anatomical Therapeutic Chemical (AUC) classification system. The ATC system divides drugs into 5 levels from anatomical group (level 1) to chemical substance (level 5). Drug targets which were classified as level 3 (therapeutic subgroup) and level 5 (specific drug) were used in this study. We used 156 GSs from level 3, and 710 gene sets from level 5 whose lengths were ≥ 5 genes from the curated GSs of Ruderfer, et al. ^30^.

To compare the current results with previous results, known EPI genes were downloaded from two sources. The first was 76 genes from https://www.cureepilepsy.org/egi/genes.asp ^11^, and the second was 218 genes from https://www.omim.org/phenotypicSeries/PS308350 of the Online Mendelian Inheritance in Man, OMIM ^71^.

#### Transcriptomic data

Gene expression specific for tissues were downloaded from the GTEx project (V6p) ^29^. We used *log*2(*x_ij_* + 1) in our analyses in which *x_ij_* was the expression value of the *i_th_* gene at the *j_th_* tissue. Spatiotemporal transcriptomic data were downloaded from BRAINSPAN ^49^. As in our previous work ^72^, this data set was partitioned into eight developmental time points (four pre-natal and four post-natal) for each of the four brain regions: the frontal cortex, temporal and parietal regions, sensory-motor regions, and subcortical regions. To create heatmaps for the main analysis, we calculated average expression across samples for each spatiotemporal point and then standardized these values. The package *mclust* ^73^ was used to cluster these standardized expression data. We also created heatmaps for each of the brain regions across 8 developmental time points. We standardized expression values across samples of the tested region and then made a heatmap for all samples.

Single-cell RNA sequencing (scRNAseq) data were obtained from Skene, et al. ^74^. Briefly, this data set included 9970 mouse cells. These cells were clustered into 24 Level 1 brain cell types and 149 Level 2 cell types ^74^. 24 Level 1 cell types were used in this study.

#### Variant data

We used DN and rare CC data of NDDs from our previous publication ^18^, a recent EPI study ^13^ and CHD data from the denovo-db database ^75^. The data of Nguyen, et al. ^18^ were collected from multiple publications and were described in detail in Table S1 of Nguyen, et al. ^18^. In summary, the DN data included 5122, 4293, 1012 and 356 trios for ASD, DD, ID and EPI respectively, 404 cases for ASD, 3654 controls ASD respectively. We also used CHD data of 1213 trios from Homsy, et al. ^28^. Variants were annotated and divided into different categories. There were categories which included loss of function (LoF) variants/mutations, missense damaging (MiD) variants/mutations. The data from Epi K. consortium and Epilepsy Phenome/Genome Project ^13^ consisted of 5696 samples: 640 cases of familial genetic generalized epilepsy, 522 cases of familial non-acquired focal epilepsy, 662 cases of sporadic non-acquired focal epilepsy and 3877 controls. We used the ultra-rare variant counts of all genes from Table S10, S11, S12 of Epi K. consortium and Epilepsy Phenome/Genome Project ^13^. These variants had minor allele frequencies <= 0.05% and MAF = 0% in ExAC (http://exac.broadinstitute.org/about) and in EVS (http://evs.gs.washington.edu/EVS/). They were annotated by SnpEff ^76^ as loss-of-function, inframe indels, or missense “probably damaging” predicted by PolyPhen-2 (HumDiv). Based on these data sets and annotations, DD, ID and CHD had only two DN categories (LoF and MiD); ASD had two DN categories (LoF, MiD) and one LoF+MiD CC population sample; EPI had two DN categories (LoF, MiD), and three CC population samples. In addition, we also used an independent EPI data set of 197 trios ^41^ to validate our results. This data set is whole-genome-sequencing (WGS) data of individuals with EPI and DD and their parents.

#### Simulated data

To evaluate the new method, ASD genetic parameters were used to simulate DN and CC data. Simulation parameters were from previous ASD studies ^10,18^ as described in Table S2. We first simulated exact parameters of ASD to compare gene counts between gTADA and extTADA and test type I errors of gTADA in the identification of eGSs. After that, we simulated different sample sizes to have a better understanding of gTADA. There were three sample sizes: case, control and family numbers. Therefore, to reduce the complexity of the simulation process, only family numbers were changed.

### Method

#### The gTADA pipeline

gTADA was designed with two main aims. The first aim is to test the enrichment of a gene set directly from DN+CC data. The second is to use enriched gene sets as prior information to improve the identification of novel significant genes associated with the tested trait—this is considered a key feature of the pipeline.

The main pipeline of gTADA is described in Figure 1 and is presented in the Results section. In summary, gTADA combined *de novo* mutations, rare inherited/case-control variants and pathway/gene-set (GS) information to jointly estimate genetic and enrichment parameters. GS information could be from gene sets or from expression data. For variant data of each gene, we used the statistical models of extTADA as described in Table S1. For GS data, there were two situations. If that was a gene set, we coded a gene as 1 or 0 corresponding with the presence or absence in all tested genes. If that was gene expression data, we used log2(1 + expression values). To incorporate GS information, we improved the main approach of extTADA. We assumed that for each *i^th^* gene, there was a probability *π_i_* for the gene to be a risk gene. This was connected to a GS by 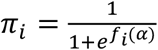 with *f_i_*(*α* = *α*_0_ + *GS* * *α*_1_ or to multiple GSs by 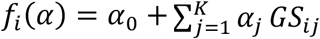. Let *X_i_* be a vector of *de novo*, case/control data of the i^th^ gene, then the likelihood (LK) function across genes was:

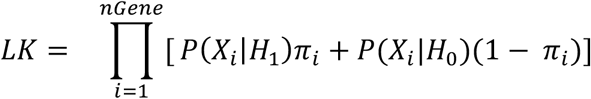

*P*(*X_i_*|*H_j_*) was the product of probabilities across all *de novo* and case/control data. If there were multiple variant categories then *P*(*X_i_*|*H_j_*) was also the product of probabilities across these categories as described in Nguyen, et al. ^18^. Table S1 describes the distribution of *de novo*, case and control data for one category of a given gene.

For a single GS model, based on the result of the equation above, a GS was considered an enriched GS (eGS) if the low boundary of its credible interval (CI) was positive. We did not adjust gene lengths inside the GS model because the statistical models of *de novo* data adjusted mutation rates (Table S1) and mutation rates were positively correlated with gene lengths.

For a multiple GS model, from eGSs, we chose a group of optimal gene sets that improved the model fit. We started with the model without any gene set (only *α*_0_). Then, we looped over all gene sets, and a gene set was added into the model if it improved the value of the likelihood function by a given threshold and the 95% CI was positive. To reduce a computational burden, we used a reduced forward-selection strategy. All enriched GSs were sorted ascendingly according to their corresponding *α* values, and GSs were added into the combined model based on this order. The final optimal gene sets were used in the identification process of risk genes. Their *α* values and genetic parameters were re-estimated to use for the calculation of posterior probabilities (PPs).

#### Generation of simulated data

To evaluate gTADA, we simulated the data as follows:

1. Simulate data without GSs:

– Input *α*_0_ to calculate 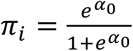 for the i^th^ gene.
– Sample the characteristics of a gene (risk or not-risk genes) *z_i_* ~ *Binomial*(2, *π_i_*):

- *z_i_* = 1 (*risk gene*): *γ_i_* ~ *Gamma*(*γ̄* * *β,β*)
- *z_i_* = 0 (*not* – *risk gene*): *γ_i_* = 1.
– Sample CC and DN counts for each gene from statistical models in Table S1.
2. Simulate gene sets:
We simulated different GS sizes. To simulate non eGSs, random genes were chosen from all genes. To simulate enriched GSs, we used prior information from Nguyen, et al. ^18^ as follows. Overlaps between eGSs and top significant genes from DN and CC data are not random. Therefore, to make the distribution of genes in gene sets more realistic in the simulation process, we used results from 186 candidate gene sets of our previous study for ASD. Briefly, eGSs from the 186 gene sets were chosen. We used extTADA ^18^ to obtain posterior probabilities (PPs) for genes from the simulation data, and then ranked the genes according to their PPs. After that, for each gene set, we made a table of overlapping-gene numbers between the gene set and genes in different groups (e.g., top 50 genes, 51st to 100th genes, ‥). In the simulation process, we allocated genes into different groups using this table. The allocation was also based on the gene size of each simulated gene set.

#### Estimation of genetic and gene-set parameters

We used Markov Chain Monte Carlo (MCMC) methods implemented in the *rstan* package ^77^ to jointly estimate all genetic and gene-set parameters. The convergence of each parameter from MCMC results was diagnosed by the estimated potential scale reduction statistic (*R̂*) inside the *rstan* package. The *Locfit* ^78^ was used to obtain credible intervals (CIs), modes of parameters.

To obtain eGSs for the process of risk-gene prioritization, we only used GSs whose low boundaries of CIs were positive. To obtain comparable results with other studies, we used posterior sampling results. A one-tail p value for each GS was calculated as the probability of GS’s alpha less than 0 if alpha’s posterior mode was positive and larger than 0 if alpha’s posterior mode was negative. All p values were adjusted by using the method of Benjamini and Hochberg ^25^.

#### Validation of significant genes

GeNets was used to test protein-protein interactions from the identified genes. Inside GeNets, the InWeb database which includes 428,429 pair-wise interactions involving 12,357 proteins ^42^ was used. The protein interaction set of InWeb is comprised of high confidence interactions from different databases ^43^. Connectivity p values were obtained using default parameters from the GeNets server (http://apps.broadinstitute.org/genets#computations). We also used STRING database to further obtain the information of protein interactions of genes. To test the enrichment of scRNAseq data, we used the same method described in Nguyen, et al. ^18^. The information of the probabilities of LoF tolerance was downloaded from ftp://ftp.broadinstitute.org/pub/ExAC_release/release0.3.1/functional_gene_constraint/ ^33^. The list of constrained genes was downloaded from Table S4 of Samocha, et al. ^34^.

#### Simulation of data to test the false discovery rates of top prioritized EPI genes

To check the observed false discovery rates (FDRs) of the top prioritized EPI genes, we simulated data similarly to the general simulation framework above. All genetic parameters which were estimated by gTADA for one trio population sample and three case/control population samples were used (Table S13). We used all 98 enriched GSs from the 1903 GSs.

## Supporting information

## Author’s contributions

Conceived and designed the experiments: HTN, EAS. Designed the pipeline used in analysis, performed the experiments, analyzed the data and drafted the manuscript: HTN. Analyzed single-cell data: JB, DP. Contributed reagents/materials/analysis tools: HTN, AD, AC, JB, AR, WM, NGS, LMH, WW, DMR, XX, MF, SMP, KP, MV, ABS, JH, JDB, DP, XH, PFS, EAS. Wrote the paper: HTN, AD, AC, XH, EAS.

## Acknowledgements

This work is supported by NIH grant R01MH105554 to E.A.S, and by NIH grant R01MH110555 to D.P. The Sweden exome sequencing data generation and analysis are supported by the Stanley Center for Psychiatric Research and NIH grant R01 MH077139 to P.F.S. This work was supported in part through the computational resources and staff expertise provided by Scientific Computing at the Icahn School of Medicine at Mount Sinai. We are deeply grateful for the participation of all subjects contributing to this research.

## Supplementary Tables and Figures

Figure S1: The performance of gTADA in the prioritization of top genes for single gene sets (GSs). Left panel compares gene counts between extTADA and gTADA for different sample sizes. The left panel is for single gene sets in which random gene sets (rGSs) and enriched gene sets (eGSs) are presented side by side. These are gene counts with different posterior probabilities (PP) of 0.95 and 0.8. The right panel describes the correlation between PPs and observed false discovery rates (FDRs).

Figure S2: The performance of gTADA in the prioritization of top genes for multiple gene sets (mGSs). Left panel compares gene counts between extTADA and gTADA for different numbers of GSs: these are gene counts with different posterior probabilities (PP) of 0.95 and 0.8. The right panel describes the correlation between PPs and observed false discovery rates (FDRs) for mGSs.

Figure S3: Results of gene-set analyses from gTADA. The left picture shows a heatmap of z-scores (estimated modes/standard errors) of all gene sets across five disorders (autism spectrum disorder: ASD, intellectual disability: ID, developmental disorder: DD, epilepsy: EPI and congenital heart disease: CHD) while the right picture presents overlapping results of significantly enriched gene sets from the analysis of gTADA.

Figure S4: P-value correlation between gTADA and previous methods. These results are for 186 gene sets (GSs) analyzed in current study and in the previous study of our group. Left panels show correlations between gTADA and the two previous methods: permutation based method (PE) and posterior probability based method (PP). Right panels describe numbers of gene sets which are identified by three methods. PE used the top 500 genes with the smallest FDRs from extTADA to test the enrichment of the 186 GSs. PP calculated the sum of the posterior probabilities of a tested GS and compare the sum with those of random GS having the same size as the tested GS.

Figure S5: Correlation between the number of gene sets and observed false discovery rates (FDRs) by using different thresholds of maximum posterior probabilities (PPs). These are simulation results for enriched gene sets of epilepsy (EPI). The genetic parameters of de novo mutations and rare case-control variants are from the analysis of 356 trios + 5,704 cases and controls.

Figure S6: gTADA results for GTEx tissues. These are credible intervals (CIs) and modes estimated by gTADA for the tissues. Red color intervals are for enriched tissues after adjusting for multiple tests.

Figure S7: The genetic parameters of epilepsy (EPI) from de novo (DN) and rare case-control (CC) data sets. Y axes are mean relative risks (mean RRs) for two DN classes, and three CC population samples. X axes are the intercept in the logistic regression: 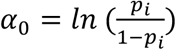, p_i_is the probability of a gene being a risk gene.

Figure S8: The number of overlapping genes between different gene sets and no GS (noGS) for epilepsy. These are the top epilepsy genes prioritized by using different types of gene sets: GTEx tissues, drug-class gene sets (DrugClassGS), drug-name gene sets (DrugNameGS) and 1901 gene sets (GS) collected from previous studies.

Figure S9: The enrichment results of single-cell RNA sequencing (scRNAseq) data in different communities. These results are for five communities generated by GeNets ^79^. For each community, scRNAseq data were tested for genes from gTADA only.

Figure S10: Single-cell based gene expressions across genes of Community 4. These are gene-expression levels standardized across genes for genes inside Community 4.

Figure S11: The enrichment results of the top epilepsy genes from gTADA and from the protein-protein interaction (PPI) analysis for different human cell types.

Figure S12: Results of the spatiotemporal gene expression analyses for the prioritized genes of different disorders: autism spectrum disorder (ASD), congenital heart disease (CHD), intellectual disability (ID) and developmental disorder (DD). These genes have maximum posterior probabilities > 0.8 from the gTADA results of candidate gene sets.

Figure S13: Spatiotemporal gene expressions across prioritized EPI genes in 4 different regions of the human brain (the frontal cortex, temporal and parietal regions, sensory-motor regions, and subcortical regions). Each heatmap is for one region and shows 8 development stages of the human brain, and each development stage has multiple collected samples. For example, columns with a red bar are for the late prenatal stage, and there are only three samples for this stage.

Table S1: Parameters of gTADA. Statistical models for de novo (dn) and case/control (cc) data are from Nguyen, et al. ^18^. *N_dn_*, *N*_1_ *and N*_0_ *are sample sizes for families, cases and controls respectively. x_dn_, x*_1_ *and x*_0_ are de novo, case and control counts in that order at a given i^th^ gene. π_i_ is the prior probability of being a risk gene for the i^th^ gene. k is the number of gene sets. GS_ij_ is the value of the j^th^ gene set at a given i^th^ gene.

Table S2: Simulation parameters for gTADA from genetic parameters of autism spectrum disorder. These parameters were from previous studies ^10,18^.

Table S3: Type I error rates of gTADA for the identification of enriched gene sets. These results are obtained by simulating non-enriched gene sets. The last column is the percentage of gene sets having the low boundaries of credible intervals (CIs) > 0. The second and third columns describe Type I errors for two approaches: p values < alpha thresholds and low CI > 0, and p values < alpha thresholds respectively.

