## Supplementary material for "Integrative analysis of rare variants and pathway information shows convergent results between immune pathways, drug targets and epilepsy genes"

### 1. Supplementary Information

#### 1.1 Results of simulated data

We tested gTADA on simulated data which included different gene-set sizes, sample sizes as well as various combinations of gene sets.

In general, when enriched gene sets (eGSs) were used, gTADA prioritized more significant genes than extTADA (Figure S1). The number of genes increased when larger gene sets were used. Regarding different sample sizes, larger differences were observed for the posterior probability (PP) threshold > 0.8. For random gene sets, the gene-count results were nearly equal between gTADA and extTADA (Figure S1).

To see the influence of gene-set sizes on PPs, the relationship between PPs and observed false discovery rates (FDRs) were also tested. For GSs less than 1000 genes, PPs > 0.8 were nearly equal to FDRs<0.1 (Figure S1). For large GSs (> 1000 genes), FDRs were slightly larger than 0.1 when PPs were > 0.8.

We also calculated Type I error rates for calling a random GS as an enriched GS. We tested for two situations: p values < alpha thresholds and CIs > 0, and only p values < alpha thresholds. gTADA had a small inflated type I error at low $\alpha$ levels, but it correctly exhibited for medium and high $\alpha$ levels. When only p values were used to test GS enrichment, the errors were slightly higher than those of both p-value and CI information, but it was still well calibrated (Table S3). Only 1.6% of random GSs were called enriched GSs (Table S3).

We also tested whether multiple GSs could increase the power of prioritizing genes for gTADA. We added GSs by using a forward-selection based strategy (Details in Methods). As expected, the number of prioritized genes increased when GS numbers increased; however, observed false discovery rates also increased (Figure S2).

#### 1.2 Results of real data.

##### 1.2.1 Comparing the results of gene-set enrichment

To test the performance of gTADA in the identification of enriched gene sets, we compared its results with our previous results (Nguyen, et al., 2017) on a set of 186 gene sets whose p values were available from the analysis of a permutation based method and a PP-based method. To compare directly with the previous results, we calculated and adjusted p values for the 186 GSs. We observed high correlations between gTADA adjusted p values and our previous methods (rho > 0.75, p ~ 0, Figure S4). gTADA was able to re-call 100% of significant gene sets identified by the permutation method, and > 89% seGS reported by the PP-based method (Figure S4).

##### 1.2.2 Calculating observed false discovery rates for different posterior probability thresholds

We performed simulation to assess observed false discovery rates (oFDRs) for different thresholds of posterior probabilities when top prioritized genes from the results of multiple gene sets. We simulated for epilepsy (EPI) because we tested this disorder in depth. The genetic parameters of EPI and enriched GSs from 1901 GSs were used in the simulation process. oFDRs increased when GS numbers increased (Figure S5). For PP > 0.95, FDRs were always less than 0.1. Similarly, FDRs were less than 0.3 with PP > 0.8.

##### 1.2.3 Insight of the rare variant genetic architecture of EPI

We applied gTADA without gene sets to infer genetic parameters of EPI (Figure S7). The mean relative risks (MeanRRs) of de novo mutations were approximately 17. For case/control (CC) data, MeanRRs of familial non-acquired focal epilepsy (familiar NAFE) and familial genetic generalized epilepsy, (familiar GGE) were nearly equal (5.0 and 4.5 respectively). Surprisingly, the mean RRs of the sporadic non-acquired focal epilepsy (NAFE) CC sample was 4.1 which was not much smaller than those of the two other CC population samples. These CC results were much larger than the result of the Epi K. consortium and Epilepsy Phenome/Genome Project (2017). Therefore, we recalculated the CC ratios and the confident intervals of these ratios for the three population samples with different PP thresholds by resampling with replacement for genes with different PP thresholds. Overall, for PP>0.1, the CC ratios were larger than 4 (Table S14). These ratios were strongly significant when we compared with random GSs having the same size from the whole genes ($p < 9.9e-4$, Table S14). Therefore, all the genes with PP > 0.1 from gTADA without GS (2102 genes) could be considered as an enriched GS across three CC population samples. This supplied more information for EPI because significant differences between cases and controls were only reported for two familiar NAFE and GGE population samples in the study of Epi K. consortium and Epilepsy Phenome/Genome Project (2017).

To better understand the results of gene sets from gTADA, we also used the same method above to test for five gene sets (43 known EPI genes; FMRP, NMDAR, seizures and ion gene sets) used in the study of Epi K. consortium and Epilepsy Phenome/Genome Project (2017). Our results were similar to the Epi K. consortium and Epilepsy Phenome/Genome Project (2017) for the four gene sets. For 43 known EPI genes, highly significant results were observed ($p < 9.9$-e4) for familiar NAFE and familiar GGE samples, but the result for sporadic NAFE sample was not significant ($p \sim 0.18$). For four other gene sets, the most significant result was the ion gene set ($p < 6.9$-e3) in the familiar NAFE samples (Table S14).

#### 2. Supplementary figures

*Figure S1: The performance of gTADA in the prioritization of top genes for single gene sets (GSs). Left panel compares gene counts between extTADA and gTADA for different sample sizes. The left panel is for single gene sets in which random gene sets (rGSs) and enriched gene sets (eGSs) are presented side by side. These are gene counts with different posterior probabilities (PP) of 0.95 and 0.8. The right panel describes the correlation between PPs and observed false discovery rates (FDRs).*

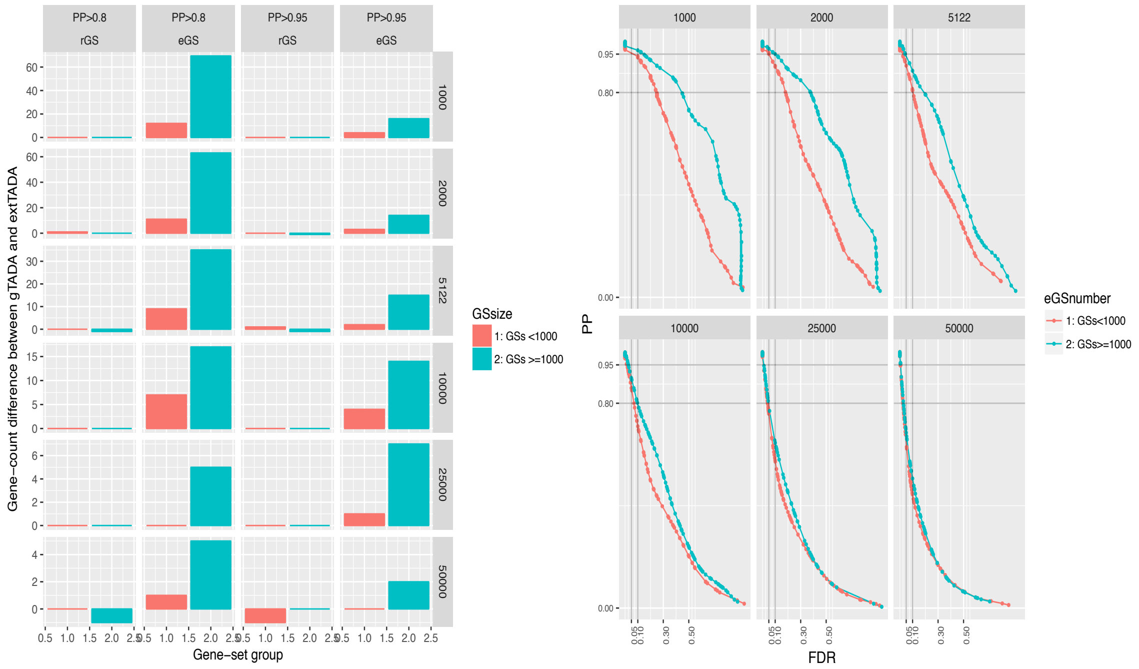

*Figure S2: The performance of gTADA in the prioritization of top genes for multiple gene sets (mGSs). Left panel compares gene counts between extTADA and gTADA for different numbers of GSs: these are gene counts with different posterior probabilities (PP) of 0.95 and 0.8. The right panel describes the correlation between PPs and observed false discovery rates (FDRs) for mGSs.*

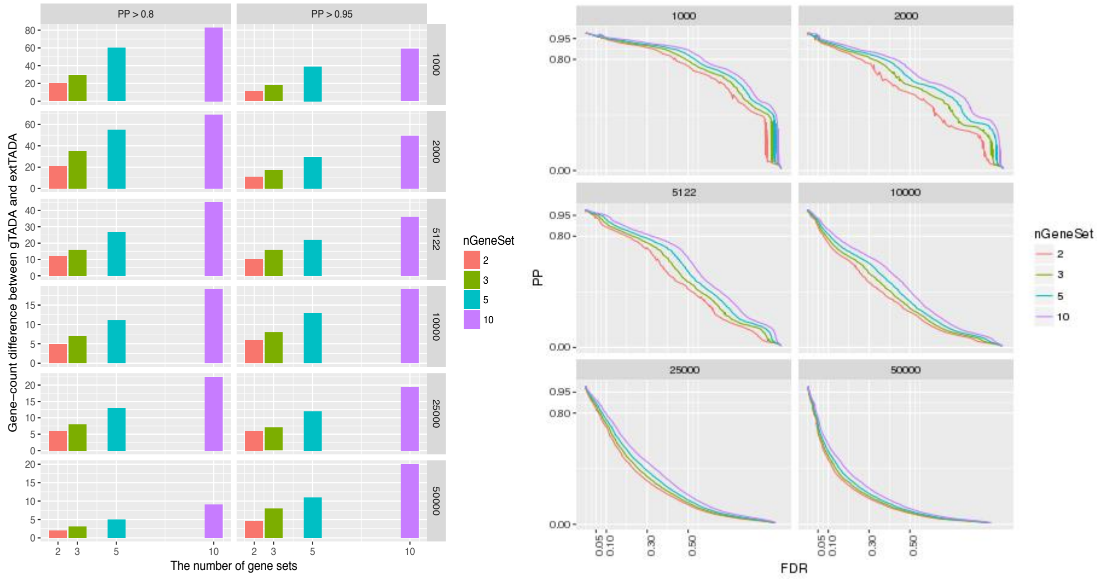

*Figure S3: Results of gene-set analyses from gTADA. The left picture shows a heatmap of z-scores (estimated modes/standard errors) of all gene sets across five disorders (autism spectrum disorder: ASD, intellectual disability: ID, developmental disorder: DD, epilepsy: EPI and congenital heart disease: CHD) while the right picture presents overlapping results of significantly enriched gene sets from the analysis of gTADA.*

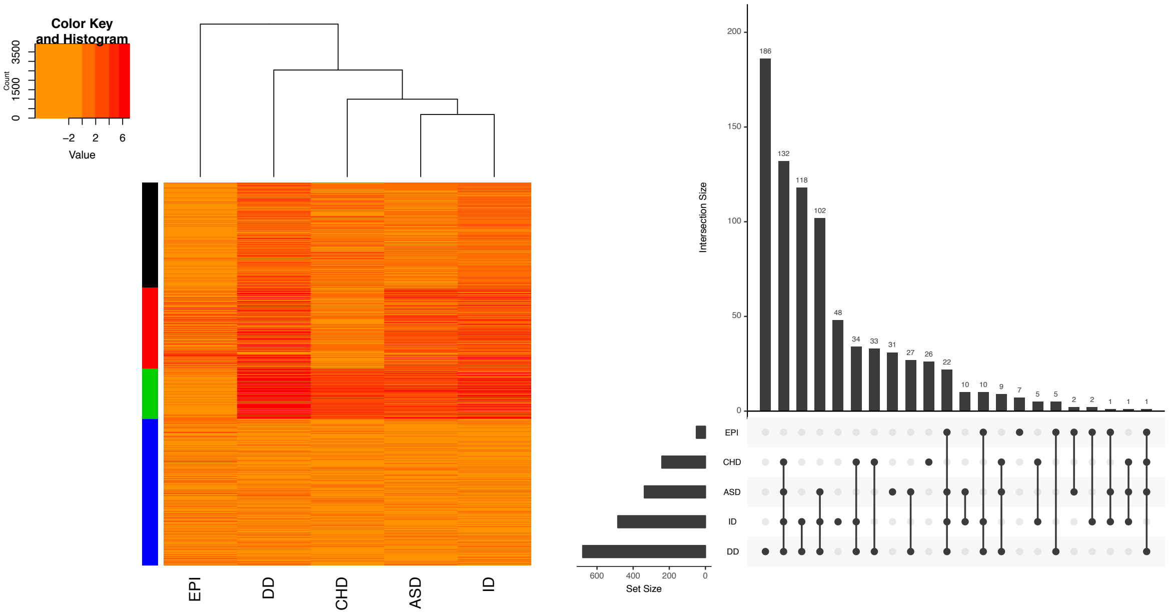

*Figure S4: P-value correlation between gTADA and previous methods. These results are for 186 gene sets (GSs) analyzed in current study and in the previous study of our group. Left panels show correlations between gTADA and the two previous methods: permutation based method (PE) and posterior probability based method (PP). Right panels describe numbers of gene sets which are identified by three methods. PE used the top 500 genes with the smallest FDRs from extTADA to test the enrichment of the 186 GSs. PP calculated the sum of the posterior probabilities of a tested GS and compare the sum with those of random GS having the same size as the tested GS.*

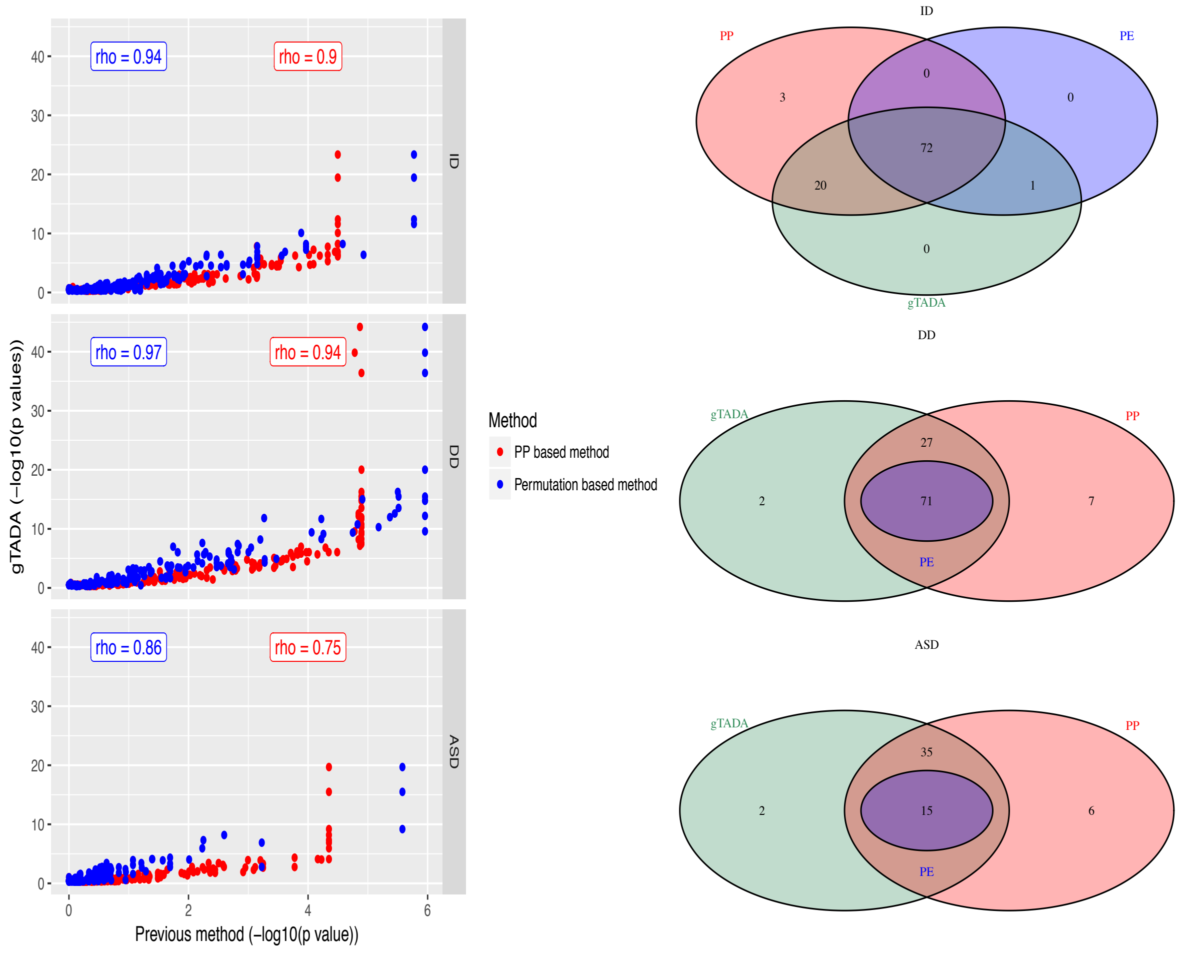

*Figure S5: Correlation between the number of gene sets and observed false discovery rates (FDRs) by using different thresholds of maximum posterior probabilities (PPs). These are simulation results for enriched gene sets of epilepsy (EPI). The genetic parameters of de novo mutations and rare case-control variants are from the analysis of 356 trios + 5,704 cases and controls.*

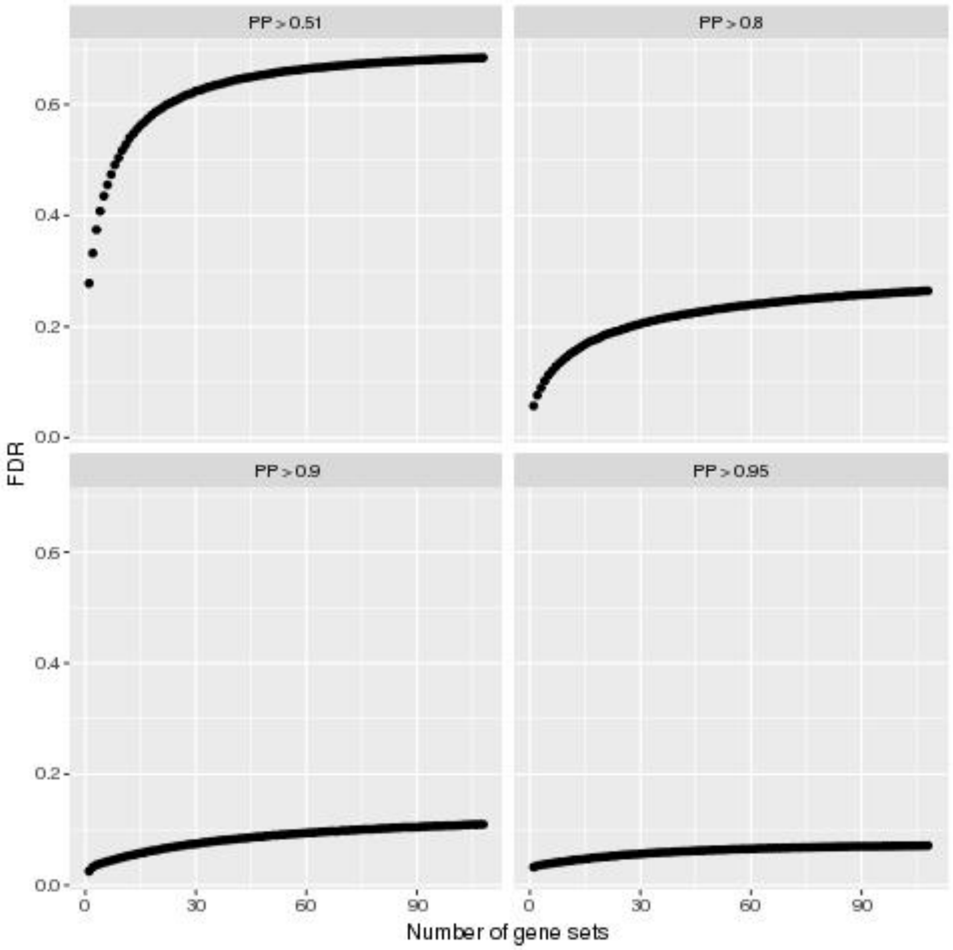

*Figure S6: gTADA results for GTEx tissues. These are credible intervals (CIs) and modes estimated by gTADA for the tissues. Red color intervals are for enriched tissues after adjusting for multiple tests.*

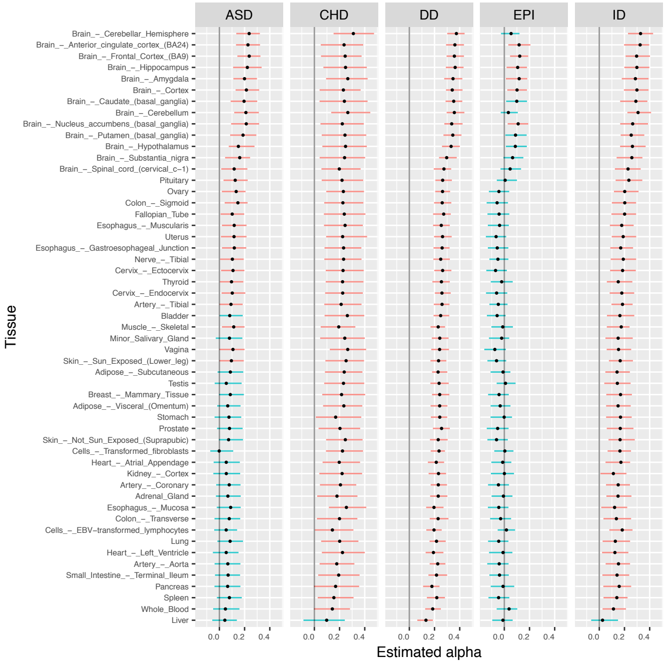

*Figure S7: The genetic parameters of epilepsy (EPI) from de novo (DN) and rare case-control (CC) data sets. Y axes are mean relative risks (mean RRs) for two DN classes, and three CC population samples. X axes are the intercept in the logistic regression:* $\alpha_0 = ln\,(\frac{p_i}{1-p_i})$*,* $p_i$ *is the probability of a gene being a risk gene.*

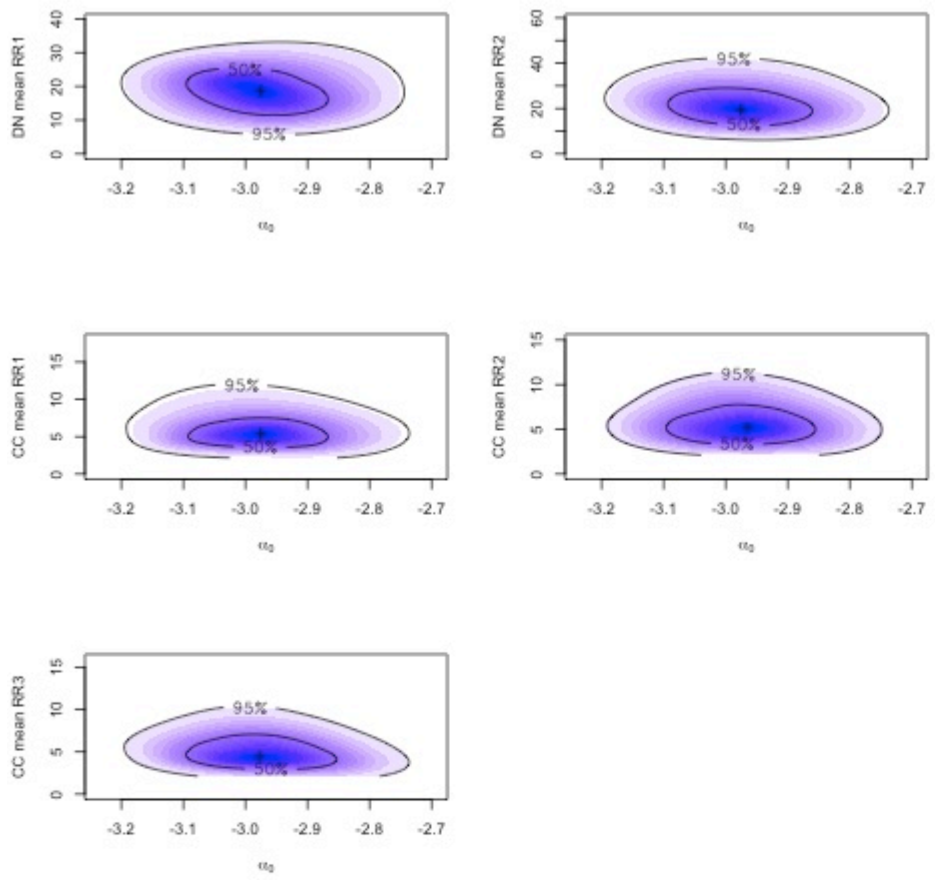

*Figure S8: The number of overlapping genes between different gene sets and no GS (noGS) for epilepsy. These are the top epilepsy genes prioritized by using different types of gene sets: GTEx tissues, drug-class gene sets (DrugClassGS), drug-name gene sets (DrugNameGS) and 1901 gene sets (GS) collected from previous studies.*

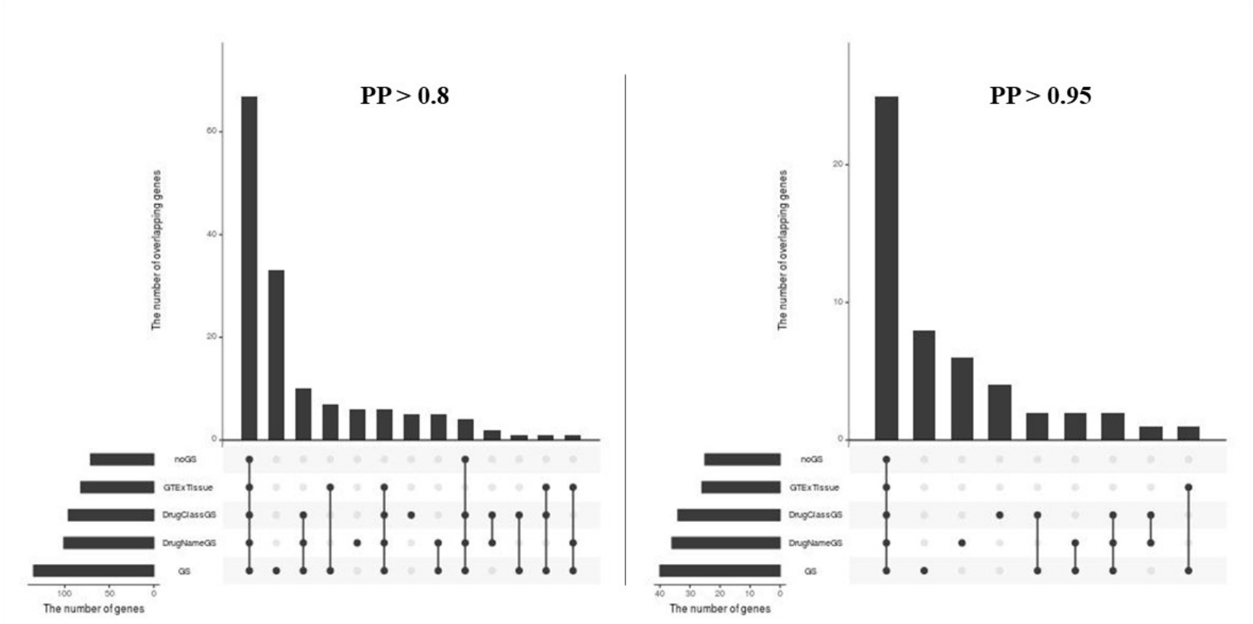


*Figure S9: The enrichment results of single-cell RNA sequencing (scRNAseq) data in different communities. These results are for five communities generated by GeNets (Hu, et al., 2017). For each community, scRNAseq data were tested for genes from gTADA only.*

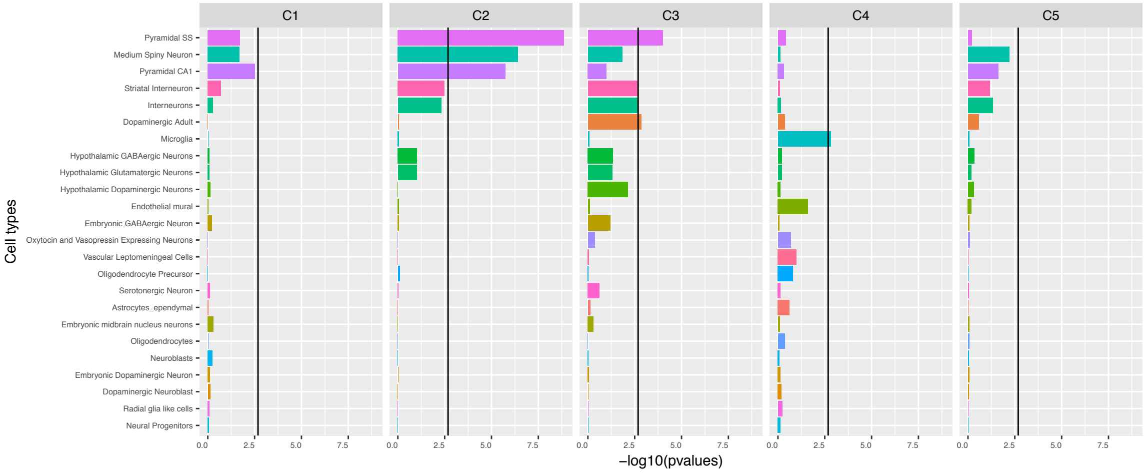

*Figure S10: Single-cell based gene expressions across genes of Community 4. These are gene-expression levels standardized across genes for genes inside Community 4.*

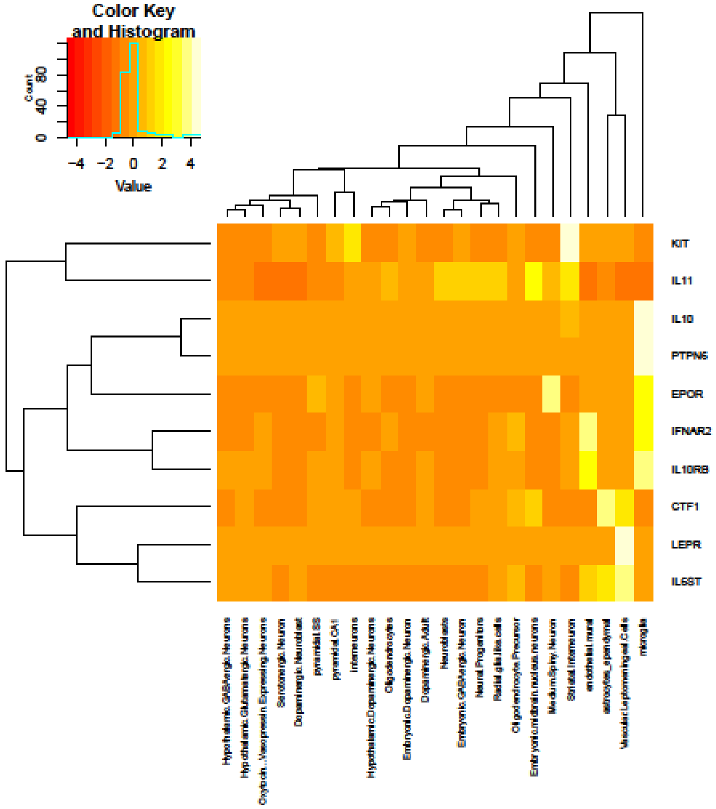

*Figure S11: The enrichment results of the top epilepsy genes from gTADA and from the protein-protein interaction (PPI) analysis for different human cell types.*

**Cell-type enrichments**

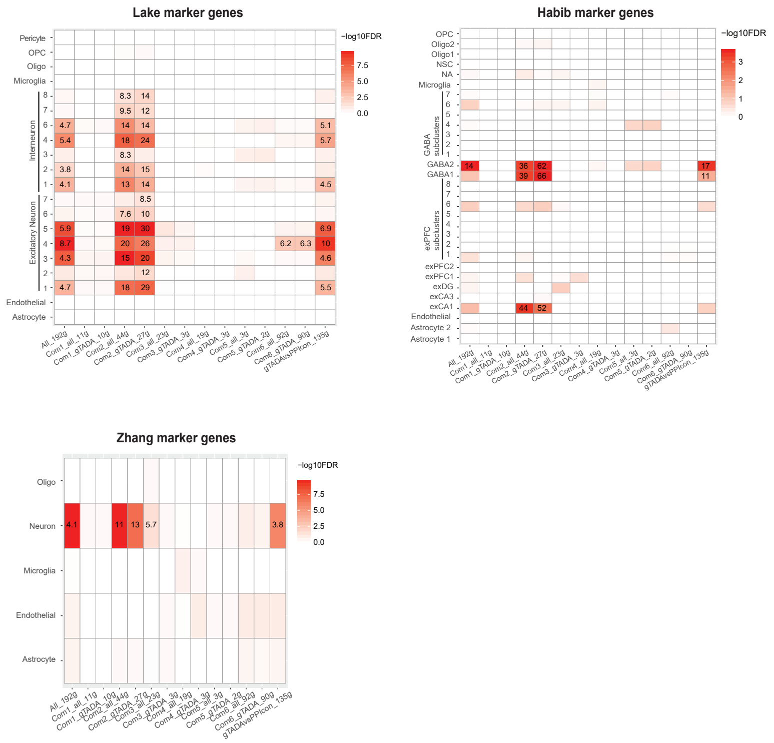

*Figure S12: Results of the spatiotemporal gene expression analyses for the prioritized genes of different disorders: autism spectrum disorder (ASD), congenital heart disease (CHD), intellectual disability (ID) and developmental disorder (DD). These genes have maximum posterior probabilities > 0.8 from the gTADA results of candidate gene sets.*

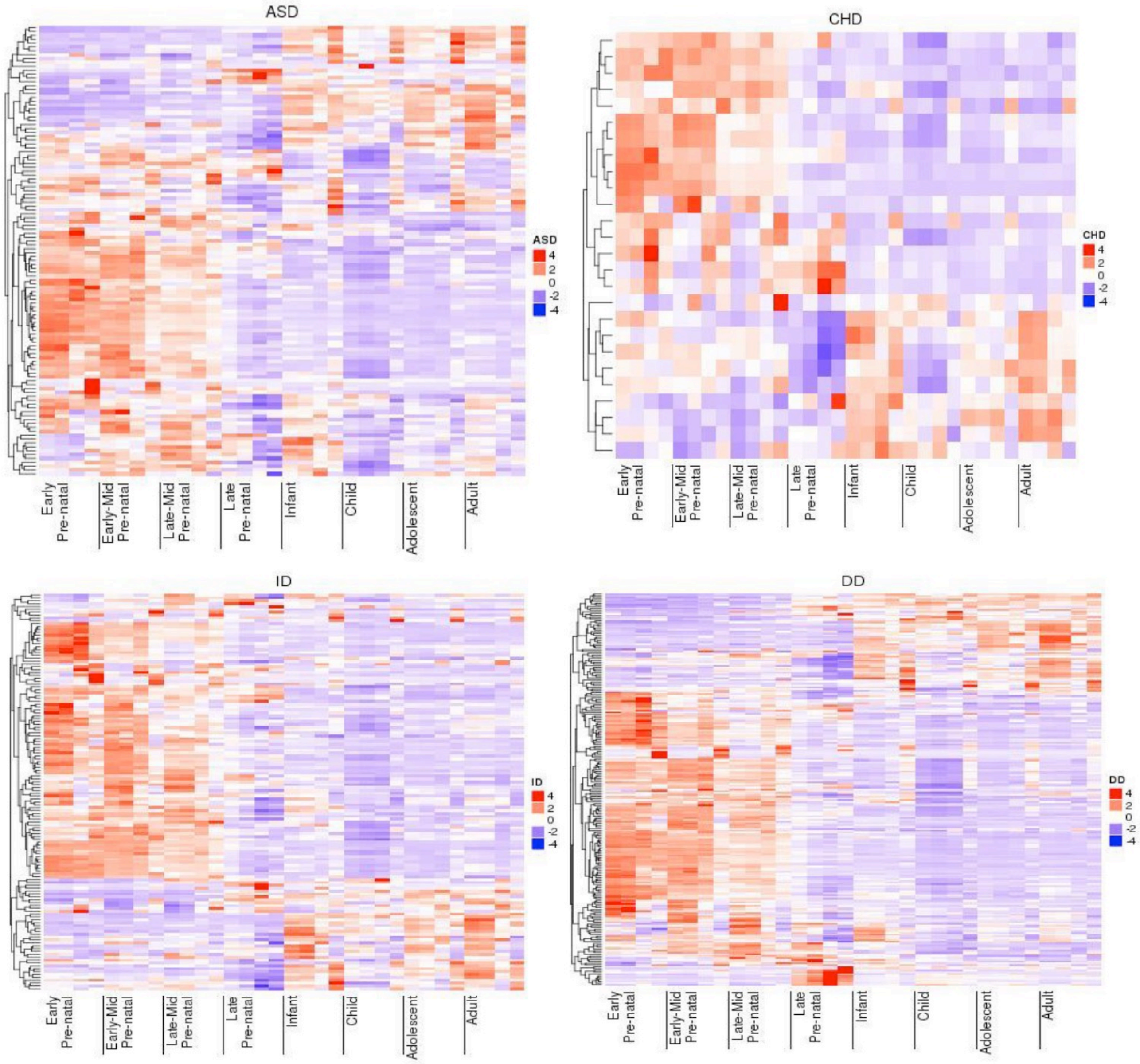

*Figure S13: Spatiotemporal gene expressions across prioritized EPI genes in 4 different regions of the human brain (the frontal cortex, temporal and parietal regions, sensory-motor regions, and subcortical regions). Each heatmap is for one region and shows 8 development stages of the human brain, and each development stage has multiple collected samples. For example, columns with a red bar are for the late prenatal stage, and there are only three samples for this stage.*

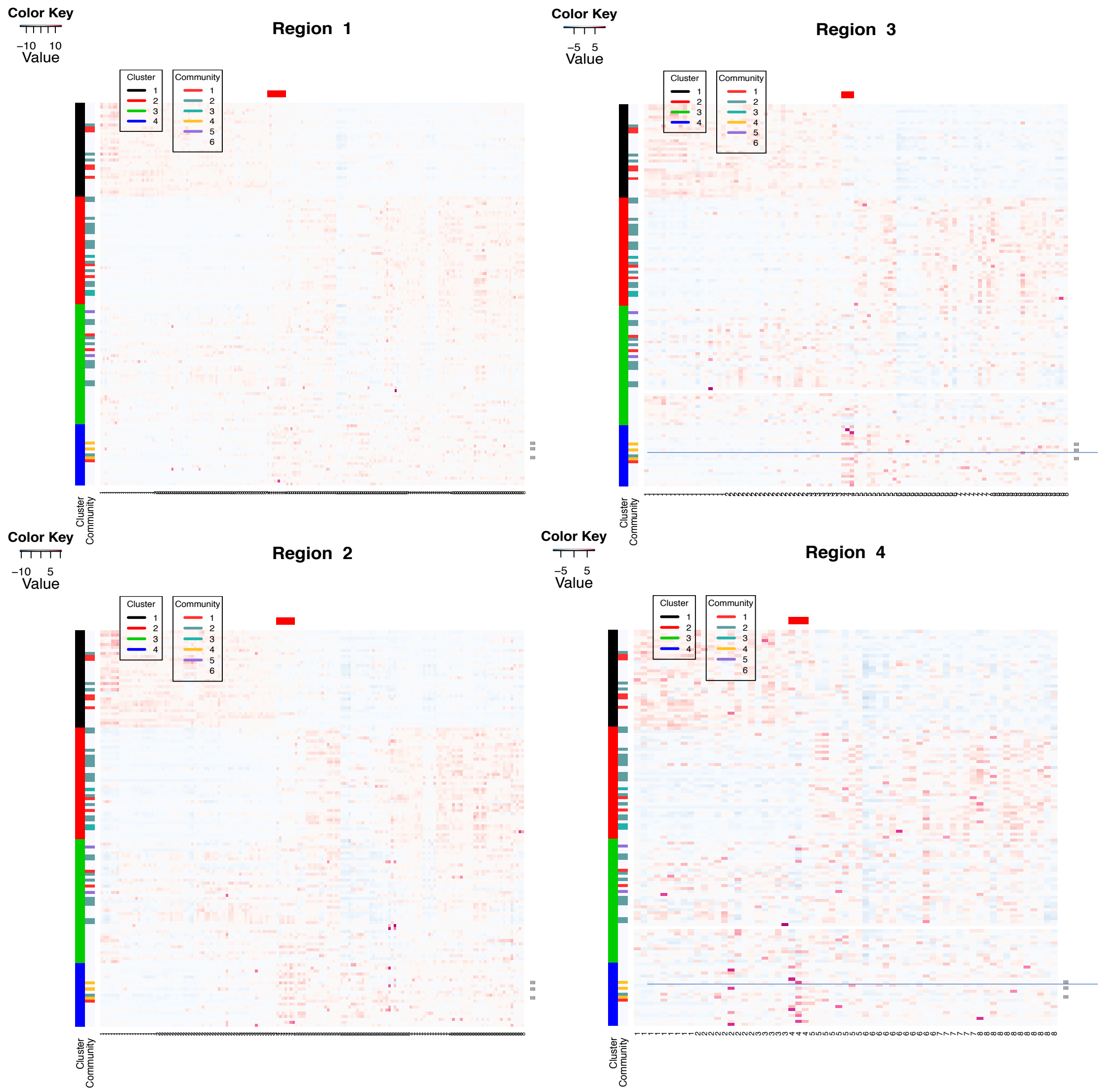

#### 3. Supplementary Tables

*Table S1: Parameters of gTADA. Statistical models for de novo (dn) and case/control (cc) data are from (Nguyen, et al., 2017). $N_{dn}, N_1$ and $N_0$ are sample sizes for families, cases and controls respectively. $x_{dn}, x_1$ and $x_0$ are de novo, case and control counts in that order at a given i^th gene. $\pi_i$ is the prior probability of being a risk gene for the $i^{th}$ gene. K is the number of gene sets. $GS_{ij}$ is the value of the $j^{th}$ gene set at the given $i^{th}$ gene.*

| Data model/Equation | Parameter prior | Hyper prior |
|---|---|---|
| $x_{dn} \sim Poisson(2N_{dn}\mu\gamma_{dn})$ | $\gamma_{dn} \sim Gamma(\bar{\gamma}_{dn} * \beta_{dn}, \beta_{dn})$<br>$\beta_{dn} = e^{a*\bar{\gamma}_{dn}^{b}+c}$ | $\bar{\gamma}_{dn} \sim Gamma(\bar{\bar{\gamma}}_{dn}, \bar{\beta})$ |
| $x_{ca} \sim Poisson(2N_1 q\gamma_{cc})$ | $\gamma_{cc} \sim Gamma(\bar{\gamma}_{cc} * \beta_{cc}, \beta_{cc})$<br>$\beta_{cc} = e^{a*\bar{\gamma}_{cc}^{b}+c}$<br>$q \sim Gamma(\rho, \nu)$ | $\bar{\gamma}_{cc} \sim Gamma(\bar{\bar{\gamma}}_{cc}, \bar{\beta}_{cc})$<br>$\frac{\rho}{\nu} = mean(\sum(x_{cn} + x_{ca}))$<br>$\nu = 200$ |
| $x_{cn} \sim Poisson(2N_0 q)$ | $q \sim Gamma(\rho, \nu)$ | $\frac{\rho}{\nu} = mean(\sum(x_{cn} + x_{ca}))$<br>$\nu = 200$ |
| $\pi_i = \frac{e^{\alpha_0 + \sum_{j=1}^{K}\alpha_j GS_{ij}}}{1 + e^{\alpha_0 + \sum_{j=j}^{K}\alpha_j GS_{ij}}}$ | $\alpha_j \sim Normal(0, 2)$ | |

*Table S2: Simulation parameters for gTADA from genetic parameters of autism spectrum disorder. These parameters were from previous studies (De Rubeis, et al., 2014; Nguyen, et al., 2017).*

| | Main parameter | Other values of parameters to test gTADA power |
|---|---|---|
| Trio numbers | 5122 | 1000, 2000, 10000, 25000 |
| Case numbers | 404 | |
| Control numbers | 3654 | |
| DN mean relative risk 1 ($\bar{\gamma}_{dn1}$) | 24.6 | |
| DN mean relative risk 2 ($\bar{\gamma}_{dn2}$) | 3.71 | |
| CC mean relative risk ($\bar{\gamma}_{cc}$) | 4.44 | |
| $\alpha_0$ | -3.1 (~ 835 risk genes) | |
| $\rho_{cc}$ | 0.66 | |
| $\nu_{cc}$ | 1947 | |

*Table S3: Type I error rates of gTADA for the identification of enriched gene sets. These results are obtained by simulating non-enriched gene sets. The last column is the percentage of gene sets having the low boundaries of credible intervals (CIs) > 0. The second and third columns describe Type I errors for two approaches: p values < alpha thresholds and low CI > 0, and p values < alpha thresholds respectively.*

| Alpha level | Type I error rate | | |
|---|---|---|---|
| | Low CI > 0 and p value < alpha | P value < alpha | Low CI > 0 (%) |
| 1.00E-04 | 5.51E-04 | 5.51E-04 | 1.6 |
| 2.00E-04 | 9.52E-04 | 9.52E-04 | 1.6 |
| 5.00E-04 | 1.70E-03 | 1.70E-03 | 1.6 |
| 1.00E-03 | 2.90E-03 | 2.90E-03 | 1.6 |
| 1.00E-02 | 1.22E-02 | 1.43E-02 | 1.6 |
| 2.00E-02 | 1.49E-02 | 2.47E-02 | 1.6 |
| 2.50E-02 | 1.53E-02 | 2.92E-02 | 1.6 |
| 3.00E-02 | 1.55E-02 | 3.43E-02 | 1.6 |
| 5.00E-02 | 1.58E-02 | 5.55E-02 | 1.6 |

#### ***Other tables are in SupTables (*****SupTable_gTADA.xlsx, SupData)**

| | | Sup Table Name | Sheet Name |
|---|---|---|---|
| Table | S4 | All gene sets used in this study | FullGeneSet |
| | S5 | Gene-set (GS) results from gTADA | GeneSetResults |
| | S6 | Prioritized genes for all disorders from gTADA (based on gene sets) | pGenesFromGSs |
| | S7 | GTEx-tissue results from gTADA | GTExResults |
| | S8 | Prioritized genes for all disorders from gTADA (based on GTEx tissues) | pGenesFromGTEx |
| | S9 | Drug-target gene-set results from gTADA | DrugTargetResults |
| | S10 | Drug-class gene-set results from gTADA | DrugTargetResults |
| | S11 | Prioritized genes for all disorders from gTADA (based on drug-name gene sets | pGenesFromDrugTarget |
| | S12 | Prioritized genes for all disorders from gTADA (based on drug-class gene sets) | pGenesFromDrugClass |
| | S13 | Genetic parameters of EPI in de novo + case/control model. | EPI_geneticPars |
| | S14 | Case/control ratios of EPI population samples. | EPI_CCratioUseBootstraping |
| | S15 | GeNets enrichment results | GeNetsEnrichment |
| | S16 | De novo counts and case/control ratios for clusters from gene expression analyses | CountsFromClustersCommunities |
